# TCR/CD3-based synthetic antigen receptors (TCC) convey superior antigen sensitivity combined with high fidelity of activation

**DOI:** 10.1101/2023.03.16.532775

**Authors:** Timo Peters, Vanessa Mühlgrabner, Rubí M.-H. Velasco Cárdenas, René Platzer, Janett Göhring, Benjamin Salzer, Angelika Plach, Maria Höhrhan, Iago Doel Perez, Vasco Dos Reis Goncalves, Jesús Siller Farfán, Manfred Lehner, Hannes Stockinger, Wolfgang W. Schamel, Kilian Schober, Dirk H. Busch, Michael Hudecek, Omer Dushek, Susanna Minguet, Johannes B. Huppa

## Abstract

Low antigen sensitivity and a gradual loss of effector functions limit the clinical applicability of chimeric antigen receptor (CAR)-modified T-cells and call for alternative antigen receptor designs for effective T-cell-based cancer immunotherapy. Here we applied advanced microscopy to demonstrate that TCR/CD3-based synthetic constructs (TCC) outperform second-generation CAR formats with regard to conveyed antigen sensitivities by up to a thousand-fold. TCC-based antigen recognition occurred without adverse non-specific signaling, which is typically observed in CAR-T-cells, and did not depend - unlike sensitized peptide/MHC detection by conventional T-cells - on CD4- or CD8- coreceptor engagement. TCC-endowed signaling properties may prove critical when targeting antigens in low abundance and aiming for a durable anti-cancer response.

## INTRODUCTION

Redirecting T-cell specificity towards tumor-associated antigens (TAAs) via CARs has proven an effective treatment strategy for patients with relapsed and refractory acute lymphoblastic leukemia (ALL), non-Hodgkin lymphoma or multiple myeloma (MM) ^1–5^. While long-term progression-free survival does frequently occur following CAR-T-cell therapy (CAR-T), relapsing disease is also common among treated patients, limiting overall effectiveness ^4, 6, 7^. The main two mechanisms underlying cancer reemergence after CAR-T concern tumor-intrinsic antigen escape ^2, 6, 8–10^ as caused by downregulation or by complete loss of TAA-expression, as well as long-term CAR T-cell exhaustion ^11, 12^ resulting from aggravated tonic CAR-mediated signaling.

Being equipped with a synthetic antigen receptor, CAR-T-cells require 1,000 or more TAAs for activation ^13^. This contrasts the remarkable capacity of conventional T-cells to detect via their naturally evolved T-cell antigen receptors (TCRs) the presence of even a single antigenic peptide-loaded MHC (pMHC), which are vastly outnumbered by structurally similar yet non-stimulatory bystander pMHC ^14–16^.

The biophysical and cell biological mechanisms underlying sensitized T-cell antigen recognition are not well understood, especially in view of the rather transient nature of stimulatory TCR-pMHC interactions. The latter typically feature lifetimes in the order of seconds and micromolar rather than antibody-like nanomolar affinities. The persistent gap in understanding is in large part caused by the complexities inherent to TCR-triggering and proximal signaling: Unlike receptor tyrosine kinases, the TCR is devoid of intrinsic kinase activity ^17^ and lacks - unlike CARs - cytoplasmic signaling domains. It is instead non-covalently associated with the invariant chains of the CD3 complex, namely a CD3γε- and a CD3δε-heterodimer as well as a disulfide-linked CD3ζ- homodimer ^18, 19, 20^, which contain, among other intracellular cytoplasmic motifs, so-termed immunoreceptor tyrosine-based activation motifs (ITAMs). The latter become tyrosine-phosphorylated upon canonical signaling by the TCR-proximal kinase Lck, which is also associated with the cytoplasmic tails of CD4- or CD8-coreceptors ^21^. This may explain why simultaneous extracellular coreceptor-engagement with a non-polymorphic region within the same MHC is critical for sensitized antigen detection via TCR/CD3 ^15, 16, 22^. In a pivotal subsequent step, Zeta-chain-associated protein kinase 70 (ZAP70) binds via its two SH2-domains to phosphorylated ITAMs (pITAMs). Once ITAM-associated, ZAP70 becomes activated by Lck or other already CD3-recruited and activated ZAP70-copies ^23^. Downstream signaling is highly amplified through ZAP70-mediated tyrosine-phosphorylation of adapter proteins and signaling effectors as well as co-stimulatory signaling cascades involving Tec family kinases ^24^.

CAR signaling resembles TCR-signaling in many but not all aspects which likely explains reduced sensitivity and long-term CAR-T-cell fitness. Membrane-proximal costimulatory signaling modules adopted from CD28, 4-1BB or OX40, which are entirely missing from the TCR/CD3 complex, keep the ITAM-containing CD3ζ-derived signaling module at a greater distance from the plasma membrane with conceivable consequences for Lck-accessibility. Furthermore, most of the CARs’ matching TAAs do not support co-receptor-mediated sensitization for lack of a co-receptor-binding site. In addition, the total number of ITAMs available for signaling amounts in monomeric CARs to only 30% and in dimeric CARs to 60% of that of a fully assembled plasma membrane-resident TCR/CD3 complex. Moreover, spatial ITAM arrangement differs, ITAM-diversity is reduced and other CD3 signaling motifs are entirely absent, which may affect not only efficient ZAP70-activation, but also the degree of tonic signaling and hyperactivity in the absence of antigens. In fact, the rapid recruitment of the E3-ubiquitin ligase c-Cbl to CD3ε’s intracellular proline-rich domain keeps non-specific TCR-proximal signaling in the absence of antigen at non-detectable levels ^25^. On the other hand, transient TCR-pMHC binding triggers TCR/CD3 effectively, but also allows for serial triggering of many TCRs by a single antigenic ligand. In contrast, CAR-signaling requires stable TAA binding, which limits in turn the overall number of triggered CARs to that of available TAAs.

Sidestepping such CAR-associated principle shortcomings may hence require a radical overhaul of conventional CAR-architectures. A rational approach would capitalize on the TCR/CD3 framework as a means to exploit TCR-associated and evolutionary preserved features of sensitized antigen detection and low tonic signaling for improved clinical efficacy. Following this rationale, several groups have fused TAA-targeting single-chain variable fragments (scF_V_s) by various means to the native TCR/CD3 complex. TCR/CD3-based synthetic constructs (TCC) including “T-cell antigen coupler” (TAC), “antibody-T-cell receptor” (AbTCR), “T-cell receptor fusion constructs” (TRuC) and “synthetic T-cell receptor and antigen receptor” (STAR) were all reported to enhance the antitumor response ^26–29^. Consistent with these observations, soluble bi-specific T-cell engagers, which directly link the TCR/CD3 complex to TAAs on target cells, have shown high clinical efficacy against B cell malignancies and uveal melanoma. At current, experimental therapies target colon, gastric, prostate, ovarian, lung, and pancreatic cancers ^30^.

Here we have applied advanced live-cell imaging to gauge the signaling performance of TCCs in much of its unfolding complexity at the single cell level and with single molecule resolution. For quantitative readouts, we confronted T-cells with planar glass-supported lipid bilayers (SLB), which had been functionalized in experimentally determined densities with the antigen itself as well as co-stimulatory and adhesion molecules to serve as defined surrogate target cell for antigen recognition. We found antigen detection mediated by TCCs about 1000-times elevated compared to that conveyed by second-generation CARs. The augmented antigen sensitivity of TCC-T-cells matched those of conventional T-cells, even in the absence of CD8 coreceptor-engagement, and was maintained in selected cases even towards low affinity antigens. Unlike second generation CARs, TCC designs did not leak antigen-independent downstream signaling, a prerequisite for durable T-cell functionality. Taken together, our findings highlight fundamental advantages that are associated with the use of TCCs especially when aiming for curative T-cell-based immunotherapies targeting tumor entities with low or highly heterogeneous TAA expression.

## RESULTS

### Anti-CD19 second generation CAR -modified T-cells (Tisagenlecleucel) fail to sense antigens in low abundance

To assess T-cell antigen sensitivities in a quantitative fashion, we confronted conventional cytomegalovirus (CMV)-specific or CD19-reactive CAR-T-cells with planar glass-supported lipid bilayers (SLB). The latter had been functionalized in defined numbers with antigen and accessory molecules to serve as surrogate target cell (Fig. 1 A) ^31^. SLBs featured the unsaturated lipids palmitoyl oleyl phosphatidyl choline (POPC, 98%) and also 1,2-dioleoyl-sn-glycero-3-[(N-(5-amino-1-carboxypentyl)iminodiacetic acid)succinyl] (nickel) (DGS-Ni-NTA, 2%), which provided a direct anchor to poly-histidine-tagged recombinant extracellular portions of the (i) antigen, e.g. HLA-A*0201 loaded with antigenic cytomegalovirus (CMV)-derived peptide pp65 (A2/CMV) for recognition by cognate RA14 TCR T-cells or CD19 for recognition by CD19-specific CAR T-cells, (ii) the adhesion molecule ICAM-1 and (iii) the costimulatory molecule B7-1 (**Fig. 1A, B**). Fluorescence recovery after photobleaching (FRAP)-based experiments testified to the lateral mobility of SLB-embedded proteins (85% and higher, **Fig. 1C**). The use of SLBs allowed also for Total Internal Reflection (TIRF) microscopy, which involves directing the fluorophore-exciting laser beam at the glass-SLB interface in a critical angle to give rise to total reflection. The resulting evanescent field penetrates into the sample no more than 200 nanometers keeping fluorescence backgrounds at levels, which are sufficiently low to allow for single fluorophore detection. As shown in **Fig. 1D, E** the use of SLBs enabled us to precisely finetune and quantitate the number of antigens and accessory proteins presented to T-cells.

**Figure 1:**
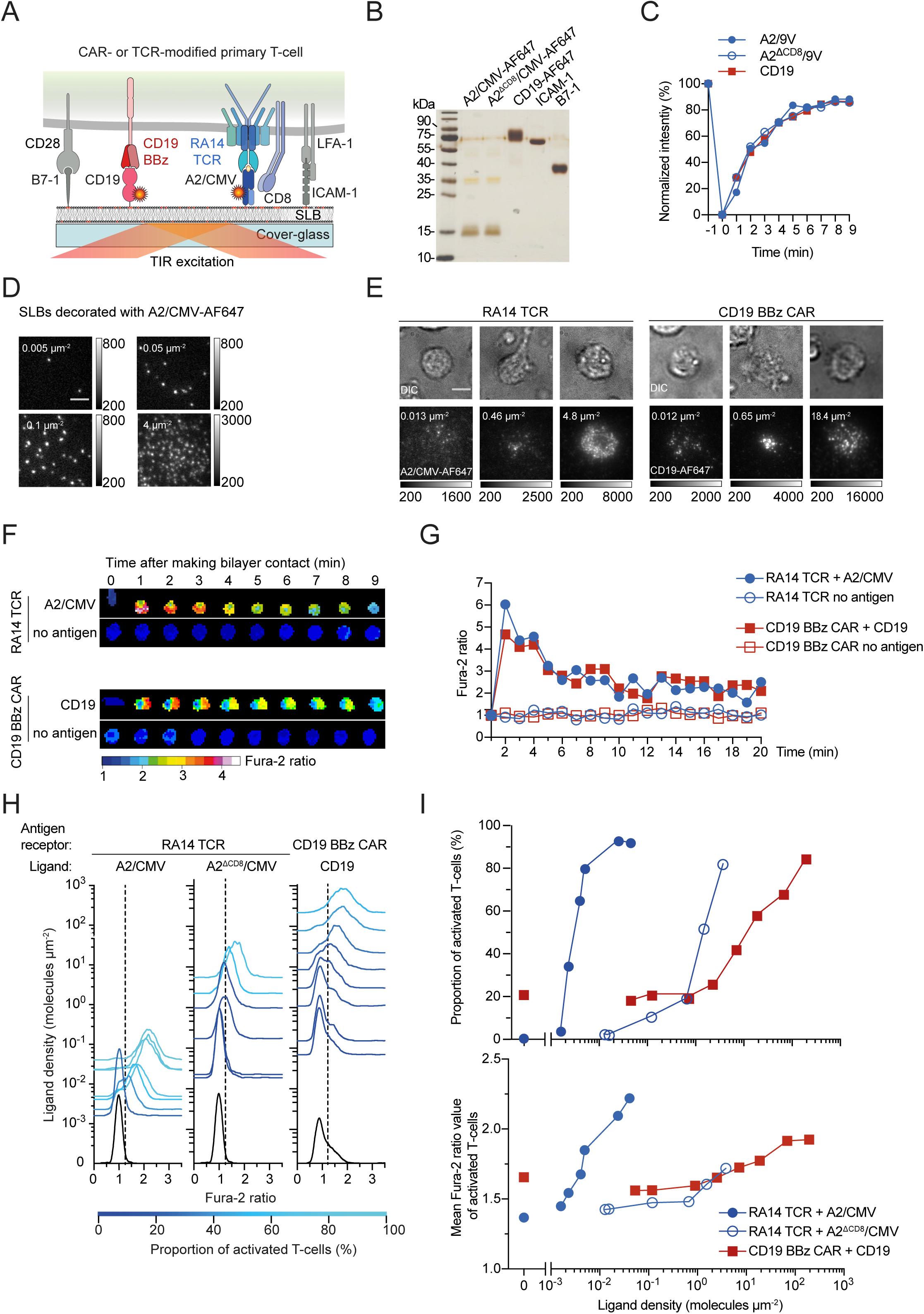
Protein-functionalized SLB-based platform for high-throughput analysis of TCR- and engineered receptor-mediated activation and downstream signaling events. (**A**) Schematic representation of an SLB equipped with fluorescently labeled antigens A2/CMV or CD19, ICAM-1 adhesion molecules and B7-1 co-stimulatory molecule for recognition by RA14 TCR T-cells or CD19 BBz CAR T-cells. Extracellular portions of proteins are extended with a polyhistidine tag (12x His) tag to interact with 18:1 DGS-NiNTA (red circles oder labeled red) present in the SLB. (**B**) 12.5% reducing SDS PAGE followed by silver staining of the recombinant proteins employed for SLB functionalization. (**C**) Fluorescence Recovery After Photobleaching (FRAP) was employed to determine the mobile fraction of SLB-anchored A2/CMV and CD19. Fluorescence intensities were normalized to the initial intensity values and plotted over time. Data shown are representative of n=3 independent experiments. (**D**) Micrographs of A2/CMV-AF647-decorated SLBs with densities indicated (white). Scale bar, 5 μm. (**E**) Micrographs showing recruitment of A2/CMV-AF647 (by RA14 TCR T-cells) and CD19-AF647 (by CD19 BBz CAR T-cells) on SLBs featuring ICAM-1 and B7-1 in addition to the respective antigen. Antigen densities indicated (white). Scale bar, 5 μm. (**F**) Antigen dependent changes in intracellular calcium concentrations as monitored by Fura-2 time-lapse microscopy in a single CD19 BBz CAR T- and a single RA14 TCR T-cell confronted with SLBs functionalized with B7-1 and ICAM-1 as well as CD19 or A2/CMV in indicated densities. (**G**) Normalized Fura-2 ratio values of T-cells shown on the left panel as a function of time. (**H**) Calcium response of RA14 TCR T-cells facing A2/CMV (left panel), RA14 TCR T-cells confronted with A2^ΔCD^^8^/CMV (middle panel) and CD19 BBz CAR T-cells exposed to CD19 (right panel) at indicated densities. Dashed lines indicate Fura-2 ratio thresholds above which T-cells were considered activated. (**I**) Upper panel: calcium response of RA14 TCR T-cells and CD19 CAR T-cells shown in (H) plotted as dose-response curves. Lower panel: mean Fura-2 ratio values of activated T-cells (population to the right of the dashed line in (H)) plotted against the employed densities of SLB-resident antigens.

For use in our experiments we isolated CD8^+^ T-cells from PBMCs of healthy donors. To confer specificity towards A2/CMV, we edited the *TRAC* locus via CRISPR-Cas9 to replace the endogenous TCR with the RA14 TCR (for a detailed description please refer to Schober K *et al*. ^32^ and the Methods section). For functional comparison, we lentivirally transduced T-cells with the FDA-approved CD19 BBz CAR (Kymriah). This CAR featured the CD19-reactive FMC63 scF_V_, a CD8α- derived hinge- and transmembrane domain, which was followed by an intracellular portion containing a 4-1BB costimulatory as well as the CD3ζ-derived activation domain (**Ext. Fig. 1B, Ext. Fig. 7, Suppl. Information**). As shown in **Fig. 1E**, both TCR- and CAR-T-cells formed stable immunological synapses with protein-functionalized SLBs and engaged their respective antigen as became visible by antigen accumulation.

To further confirm the SLBs’ functional integrity, we monitored the intracellular second messenger calcium in SLB-contacting T-cells via ratiometric live-cell imaging with the use of the calcium-sensitive dye Fura-2 AM. The rise in intracellular calcium is essential for and precedes all T-cell effector functions following antigenic stimulation. As shown in **Fig. 1F, G**, both RA14 TCR T-cells and CD19-CAR T-cells exhibited a rapid antigen-dependent increase in intracellular calcium levels shortly after contacting SLB functionalized with ICAM-1, B7-1 and optionally the corresponding antigen (A2/CMV or CD19). After reaching a peak within the first three minutes of SLB-contact, the calcium signal plateaued above the baseline levels which had been set in antigen-free control experiments.

To determine the sensitivity of RA14 TCR T-cells and CD19 BBz CAR T-cells towards antigen, we exposed T-cells to SLBs, for which we had titrated the densities of either A2/CMV or CD19, and recorded the temporal dynamics of the ensuing calcium response at the single cell level. More specifically, we tracked T-cells for each condition using a published particle tracking algorithm^24^. Tracking parameters were chosen so that on average 250 (minimally 80 and maximally 550) tracks of T-cells, which had been in contact with the SLB for at least 12 minutes, were included in the analysis. For each track we determined the mean Fura-2 ratio value of eleven 15-second intervals (2.5 minutes) following the maximum Fura-2 ratio value. Cells confronted with antigen-free SLBs served as negative control. The average of measured Fura-2 ratio values was set to 1 for the purpose of normalization. Acquired data were next organized in histograms (**Fig. 1H**) to visualize the measured distribution of normalized Fura-2 ratio values as a function of the antigen densities employed. Indicative of spurious signaling, a significant proportion of the CD19 BBz CAR- but not RA14 TCR T-cells displayed elevated Fura-2 ratio values in the absence of antigen (**Fig. 1H**, black histograms intersecting with 0 molecules µm^-2^).

As shown in **Fig. 1H, I**, the minimum density required for the activation of 34% of the RA14 TCR T-cells amounted to 0.002 molecules per μm^2^. Considering an average synaptic area of 120 µm^2^, we concluded that at least in the initial stage of contact formation each T-cell encountered on average 0.24 ligands. The fact that a sizable proportion of activated RA14 TCR T-cells had fluxed calcium almost immediately after making SLB-contact, rendered it likely that RA14 TCR-T-cells can in principle respond already to a single antigenic pMHC.

Indicative of a considerably reduced antigen sensitivity, only 42% of the CD19 BBz CAR T-cells became activated when facing CD19 at a density of 7 molecules per µm^2^. For best comparison, we plotted recorded dose response curves to calculate the ligand density required for the half-maximal response, hereafter termed the activation threshold (for details refer to Methods section). As is shown in **Table 1**, the EC50 antigen value for CD19 BBz CAR T-cell activation threshold was more than 1,000-times higher than that of RA14 TCR T-cells. Testifying to the physiological relevance of our calcium recordings, antigen levels for half maximal γ-interferon (IFN-γ) secretion were about 1,000-times higher for CD19 BBz CAR T-cells when compared to that of RA14 TCR T-cells (**Ext. Fig. 2A**).

**Table 1.**
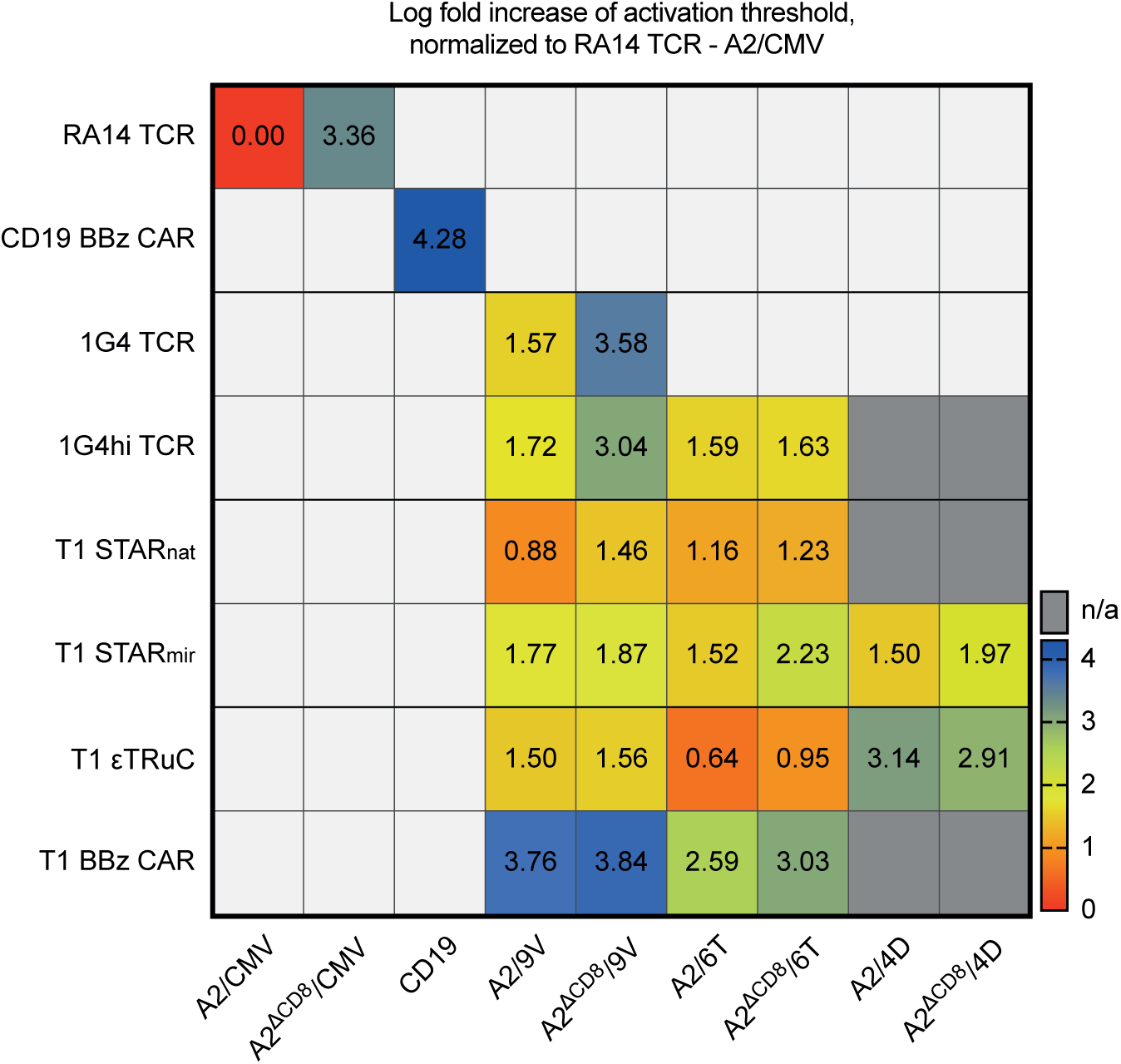

To identify reasons underlying the sizable differences in antigen sensitivities between RA14 TCR- and CD19 BBz CAR T-cells, we investigated the role of the CD8-coreceptor, which interacts with antigen-loaded MHC class I molecules but not with CD19. As pointed out above, CD8 is tethered via its cytoplasmic tail to TCR-proximal Lck, which phosphorylates upon TCR-pMHC engagement the cytoplasmic ITAMs of the CD3 chains and in a following step ITAM-recruited ZAP70 for further downstream signaling. **Fig. 1H** and **I** (upper panel) demonstrate that CD8-engagement with MHC class I was paramount to maintaining the exquisite antigen sensitivity of RA14 TCR-T-cells, as we noticed a 250-fold decrease in antigen sensitivity when stimulating T-cells with A2^ΔCD^^8^/CMV, a mutant (DT227/228KA) with abrogated CD8 binding. Once activated, RA14 TCR T-cells outcompeted CD19 BBz CAR T-cells also with regard to the amplitude of the calcium response (**Fig. 1I**, lower panel). Of note, higher levels of antigen-dependent calcium signaling in RA14 TCR T-cells were also dependent on CD8-coreceptor engagement, as abrogation of the CD8-binding-site in A2/CMV muted mean Fura-2 ratios to levels that were comparable with, if not even lower than those observed in CD19 BBz CAR T-cells.

### Surface expression of TCR/CD3-based synthetic antigen receptors

We reasoned that the therapeutic use of synthetic antigen receptors, which had been engineered along the evolved TCR/CD3 architecture, may at least in part give rise to the exquisite sensitivity and fidelity which characterize TCR-based antigen recognition. If true, such properties could in principle be exploited to not only enhance clinical efficacy but also to broaden the spectrum of permissible antigen-interaction kinetics in efforts to refine tumor specificity. However, such rationale may be derailed by the previously observed tendency of sensitized TCR-based antigen recognition to depend on CD8- or CD4-co-engagement, which is usually not supported by TAAs targeted by CARs. To address such possibility in a comprehensive manner, we dissected the contribution of CD8-mediated antigen-engagement to the recognition of an A2 restricted T-cell epitope by T-cells which had been modified with either cognate TCRs, second-generation CARs or TCCs (see below).

To be able to quantitate and directly compare the detection capacity of TCR-, CAR of TCC-modified T-cells, we chose to target the canonical cancer-testis NY-ESO-1 peptide presented in the context of HLA-A*0201 (A2) via conventional TCR-recognition or with the use of the TCR-like 3M4E5 mAb -derived T1 scF_V_, serving as the binding module for T1 CARs and T1 TCCs ^33^ (see below and **Fig. 2A**, **Ext. Fig. 3B, C**). The T1 scF_V_ engages A2/NY-ESO-1 in a peptide-specific manner, and point mutations within the NY-ESO-1 peptide allowed us to vary kinetic parameters of receptor-antigen binding and assess their role in sensitized ligand recognition (for an overview of T1-scF_V_-A2/NY-ESO-1 binding kinetics refer to **Ext. Fig. 3A** and **Table 2**). Employing A2/NY-ESO-1 as model antigen enabled us furthermore to directly compare the functional performance of synthetic antigen receptors with that of the A2/NY-ESO-specific 1G4 TCR and a genetically engineered version thereof binding A2/NY-ESO with picomolar affinity (1G4hi TCR). For direct comparison we included a second generation T1-BBz CAR as well as three different T1-TCCs.

**Figure 2:**
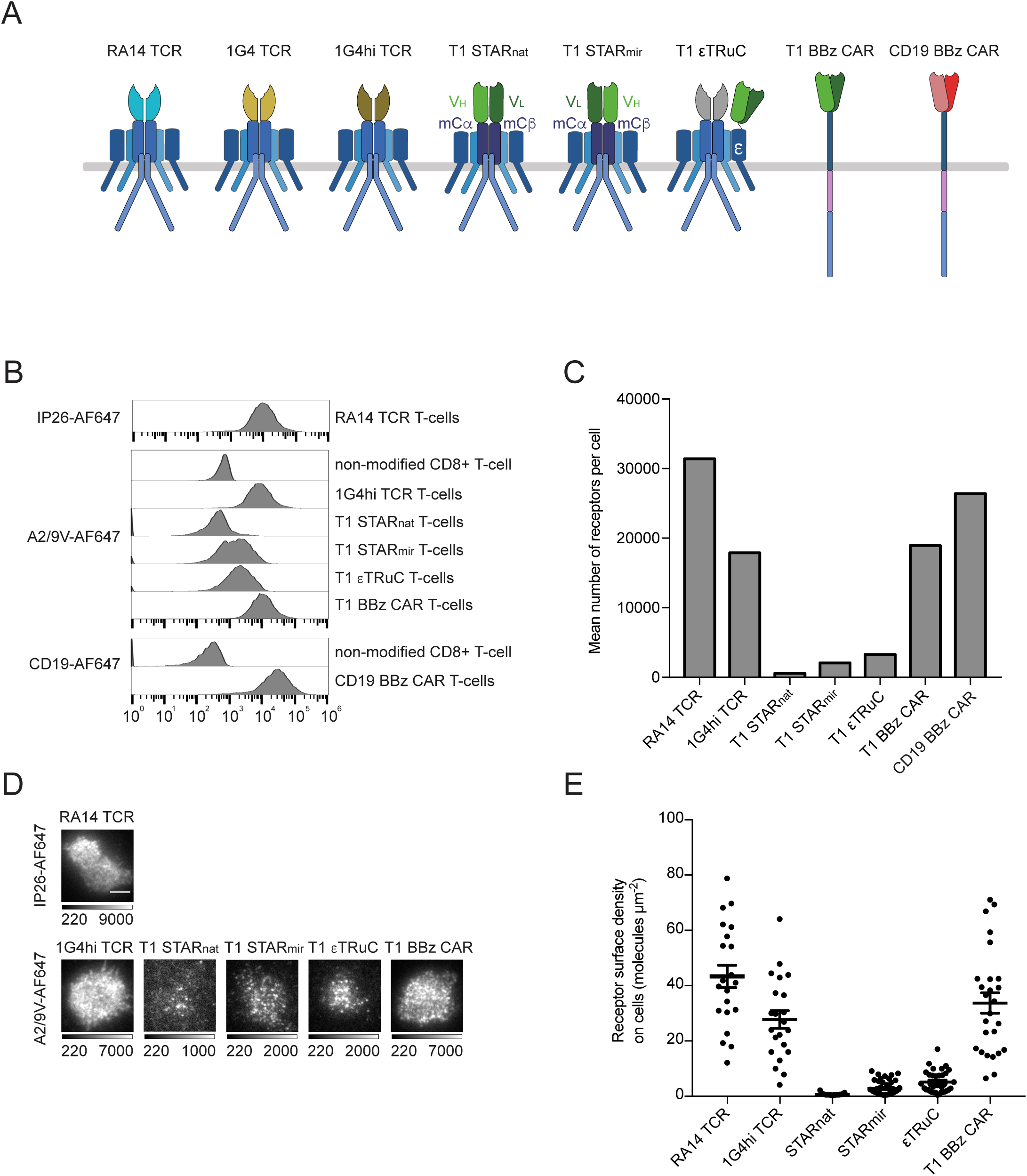
Design and surface expression of antigen receptor constructs. (**A**) Schematic representation of RA14 TCR, 1G4 TCR, affinity optimized c58c61 1G4 TCR (1G4hi TCR), synthetic dimeric receptors T1 STAR_nat_ and T1 STAR_mir_ (based on the murine TCRαβ), T1-scF_V_ tethered to CD3ε via a flexible linker (εTRuC), T1-scF_V_ BBz CAR and FMC63-scF_V_ BBz CAR. (**B**, **C)** Flow cytometric surface staining of antigen receptor modified T-cells as performed with indicated probes to determine the mean number of surface-expressed antigen receptors. Quantitation of surface receptor (C) was based on label saturation and flow cytometric calibration. (**D, E**) TIRF images of living T-cells modified with indicated antigen receptors, stained as indicated with probes under saturating conditions (for details see Ext. Fig. 1C), confronted with ICAM-1-functionalized SLBs for adhesion and quantitated for respective surface densities (n = 30 cells per condition). Error bars: sem.

**Table 2.**
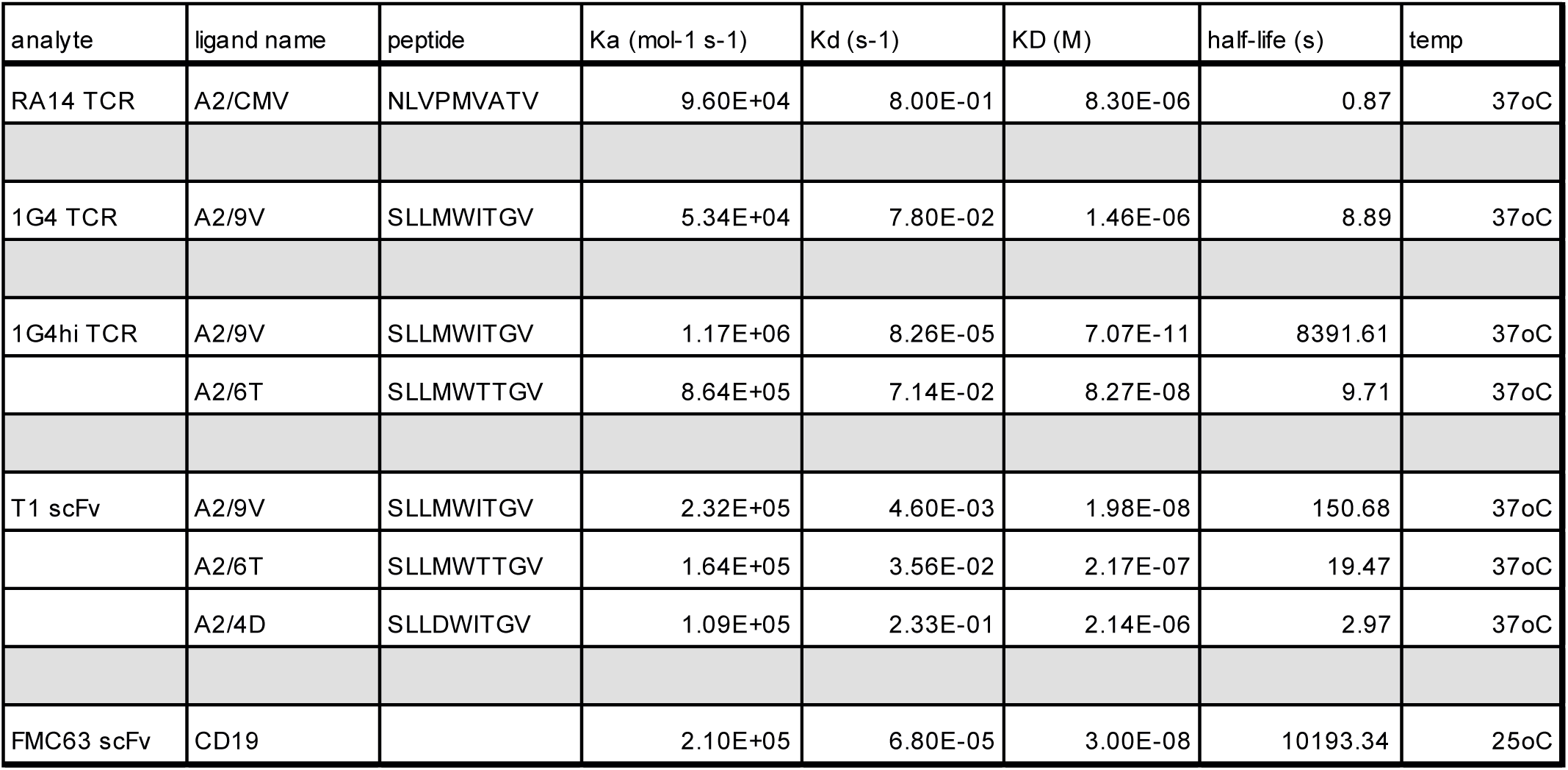

As is schematically depicted in **Fig. 2A** we constructed two <u>S</u>ynthetic <u>T</u>-cell receptor and <u>A</u>ntigen <u>R</u>eceptors (STAR), for which we replaced the V_α_- and V_β_- domains of a murine TCR with the V_L_- and V_H_- domains of the T1 scF_V_. STAR natural (T1 STAR_nat_, with V_L_ fused to Cβ and V_H_ to Cα) denotes a version which engages A2/NY-ESO-1 in a natural docking angle as is observed for most productive TCR-pMHC interactions^33^ and which preserves the orientation of CD8-binding with regard to STAR engagement. The second STAR construct termed STAR mirror (T1 STAR_mir_, with V_H_ tethered to Cβ and V_L_ to Cα) binds A2/NY-ESO-1 in the opposite orientation^33^. As an alternative to STARs, we linked the T1-scF_V_ via a flexible linker to N-terminus of CD3ε to result in a so-termed CD3ε <u>T</u>-cell <u>R</u>eceptor f<u>u</u>sion <u>C</u>onstruct (εTRuC) after its inclusion into a nascent TCR/CD3 complex.

To generate 1G4 TCR T-cells, we knocked both subunits of the 1G4 TCR into the *TRAC* locus of human primary CD8^+^ T-cells via a CRISPR-Cas9-based protocol (Ext. Fig. 1a) All other constructs were introduced by lentiviral transduction (**Fig 2B**).

Given the empiric nature of their structural design, none of the T1-based antigen receptors used in this study had undergone - unlike TCR/CD3 - natural selection with possible consequences for surface expression levels and efficacy of T-cell antigen recognition. To address this, we quantitated surface expression levels of all constructs via flow cytometry (**Fig. 2B**, **Ext. Fig. 1C, D**). The high affinity antigen interactions between FMC63- and T1 scF_V_s and their corresponding ligands allowed for quantitative surface staining via recombinant monomeric ligands, such as soluble Alexa Fluor 647 (AF647) -conjugated CD19 and A2/NY-ESO-1, respectively. The TCR-reactive monoclonal antibody (mAb) IP26 was employed to stain surface-accessible RA14 TCRs. Given picomolar affinities, 1G4hi TCR-T-cells were also quantitatively labeled with AF647- conjugated A2/NY-ESO-1. To arrive at absolute numbers of T-cell surface-resident receptors, recorded mean fluorescence intensities were calibrated with the use of Quantum^TM^ MESF (for more details, please refer to the Methods section).

As shown in **Fig 2B**, RA14 TCR was expressed in highest numbers with on average 31.000 TCRs per cell. Lentiviral expression of 1G4hi TCR, T1 BBz CAR and CD19 BBz CAR resulted in about 18,000, 19,000 and 26,000 molecules per T-cell, respectively. In contrast, surface expression of T1 STAR_nat_, T1 STAR_mir_ and T1 εTRuC amounted to only 600, 2000 and 3300 receptor entities per cell. In line with this, direct quantitation of surface receptors densities via total internal reflection microscopy (TIRF) microscopy produced the same ranking of expression: The knocked-in RA14 TCR was present at highest levels with an average of 43 TCRs per μm^2^ while T1 STAR_nat_ was barely detectable with about 60-times fewer receptors (0.68 receptors per μm^2^) (**Fig. 2D, E**).

As shown in **Ext. Fig 4A**, **B**, CRISPR-Cas9-mediated ablation of TCRα and TCRβ genes prior to lentiviral transduction of STAR constructs led to a 3- and 4.5-fold increase in surface expression for T1 STAR_nat_ and the T1 STAR_mir_, respectively. These observations were consistent with less efficient assembly of these constructs with endogenous CD3 subunits to form nascent multi-subunit complexes with CD3 during TCC biogenesis in the ER.

### T1-based TCR/CD3-based synthetic receptors confer superior sensitivity towards ligands featuring a wide range of affinities

We next assessed sensitivities conveyed by the T1 STAR_nat_, T1 STAR_mir_, T1 εTRuC and the T1 BBz CAR towards the high affinity ligand A2/9V (t_1/2, 37°C_ = 158 s, K_D_ =19.8 nM). T-cells modified with the 1G4 TCR (t_1/2, 37°C_ = 8.89 s, K_D_ =1.46 µM) and the 1G4hi TCR (t_1/2, 37°C_ = 8400 s, K_D_ =70.7 pM) served as reference for comparison. For this we confronted receptor-modified T-cells with SLBs functionalized with ICAM-1 and B7-1 and titrated densities of A2/9V, and next determined respective antigen thresholds for activation as monitored via intracellular calcium. Despite low surface antigen receptor expression levels, T1 STAR_nat_ T-cells outperformed all other constructs tested: they required a seven- or 750-times lower antigen threshold when compared to T-cells modified with the 1G4 TCR (runner up) and the trailing T1 BBz CAR, respectively (**Fig. 3**, **Ext. Fig. 5** and **Table 1**). Similarly, IFN-γ release by TCC-modified T-cells increased in an antigen density-dependent manner. In contrast, T1 BBz CAR T-cells secreted considerable IFN-γ-levels already in the absence of antigen, which rose marginally at low antigen levels and amounted to only twice to three-times the background levels when CAR-T-cells had been confronted with high antigen densities (**Ext. Fig. 2B**).

**Figure 3:**
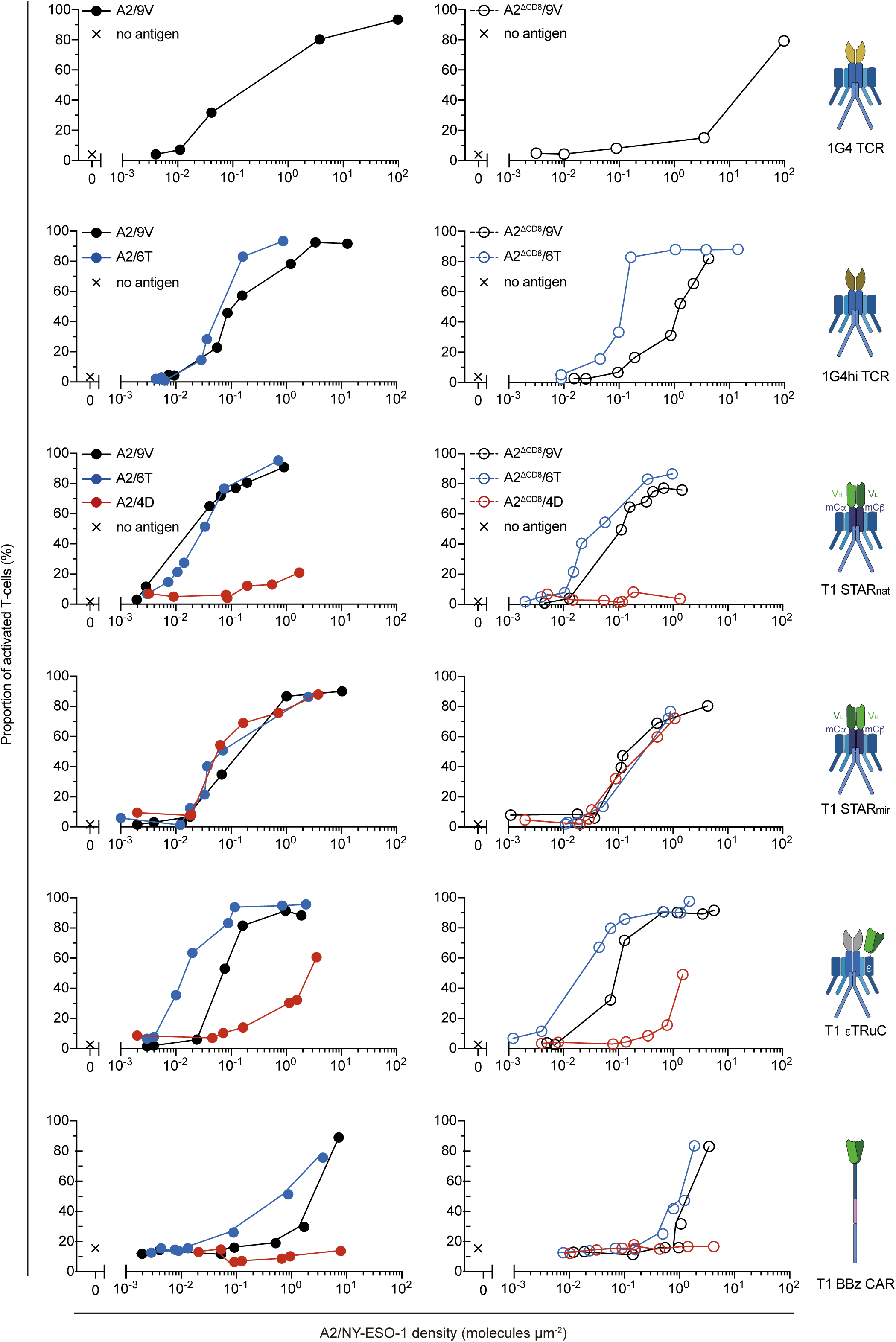
Calcium analysis of T-cells equipped with NY-ESO-1-specific TCRs, TCCs and CAR. Calcium dose-response of 1G4 TCR T-cells, 1G4hi TCR T-cells, T1 STAR_nat_ T-cells, T1 STAR_mir_ T-cells, T1 εTRuC T-cells and T1 BBz CAR T-cells which had been confronted with SLBs functionalized with A2/9V, A2/6T, A2/4D, A2^ΔCD^^8^/9V, A2^ΔCD^^8^/6T or A2^ΔCD^^8^/4D at indicated densities. Data shown are representative of (n=2) independent experiments from 2 different donors.

Indicative of a rather moderate sensitization via CD8-co-engagement, the use of A2^ΔCD^^8^/9V caused only a 5-fold reduction in sensitivity for T1 STAR_nat_ T-cells, which was slightly higher than those recorded for T1 εTRuC- and T1 STAR_mir_ T-cells. In stark contrast, T1 BBz CAR T-cells continued to lag behind in antigen detection efficiency by more than 2 orders of magnitude.

Highest T1-related sensitivities were measured for T1 εTRuC-T-cells facing the intermediate affinity A2/6T ligand (t_1/2, 37°C_ = 20 s, K_D_ = 217 nM). Such T-cells outperformed T-cells expressing the T1 STAR_nat_, the T1 STAR_mir_ or the T1 BBz CAR by a factor of 3, 17 and 90, respectively. Interestingly, both the T1 εTRuC and the T1 BBz CAR conveyed 7- and 15-times increased sensitivities towards A2/6T, when compared to those measured against the high affinity ligand A2/9V. Hence, transient rather than more stable receptor-antigen binding with off-rates resembling those of typical TCR-pMHC interactions appear to result in more productive intracellular signaling for T-cells featuring the T1 εTRuC and the T1 BBz CAR. Of note, this trend was no longer observable for T-cells expressing T1 STAR_nat_, albeit at considerably lower surface levels, which appeared to have become a limiting factor. Of note, abrogating CD8-engagement through the use of A2^ΔCD^^8^/6T barely affected the sensitivities conveyed by all four constructs. Despite a twofold loss, the T1 εTRuC still performed best, followed by the T1 STAR_nat_, the T1 STAR_mir_ and the T1 BBz CAR.

The low affinity ligand A2/4D (t_1/2, 37°C_ = 2.97 s, K_D_ =2.14 µM) was still efficiently recognized by T1 STAR_mir_ T-cells but no longer by T-cells modified with T1 STAR_nat_. Of note, a three-fold increase in T1 STAR_nat_ surface expression promoted by genetic CRISPR-Cas9-mediated ablation of endogenous TCRs did not markedly affect T-cell reactivity (**Ext. Fig. 4A, B**), rendering low T1 STAR_nat_ expression levels a less likely cause for poor recognition. In view of this, we consider it more likely that proper folding and assembly of T1 STAR_nat_ in the ER was at least in part compromised giving in turn rise to lower overall surface expression levels. Of note, T1 εTRuC T-cells showed a considerably reduced response to A2/4D when compared to T1 STAR_mir_ T-cells. This behavior may single out the rigidity of the tether between the scF_V_ and CD3ε as a critical structural parameter for the design of εTRuCs targeting low-affinity ligands. Furthermore, abrogating CD8-engagement did not majorly affect low affinity ligand detection by T-cells expressing TCR/CD3-based receptor constructs (**Fig. 3**, **Ext. Fig. 4C, D**).

In line with previous observations related to the performance of the CD19 BBz CAR, 15% of T1 BBz CAR T-cells exhibited elevated calcium signaling in the absence of antigen while none of the T-cells functionalized with TCRs or synthetic TCR/CD3-constructs showed any evidence for this behavior. Of note, we observed in essence the same signature of increased antigen sensitivity and absence of spurious activation when profiling CD19 εTRuC T-cells (**Fig. 4**). We hence conclude that TCCs maintain the fidelity of activation, a hallmark of conventional T-cells, while outperforming conventional second-generation CARs with regard to antigen sensitivity by up to three orders of magnitude. We consider it likely that reaching such degree of functionality will be critical to prevent the emergence of antigen-escape variants, i.e. by promoting sustained T-cell effector function after autologous cell transfer while sustaining exquisite T-cell sensitivity towards TAA.

**Figure 4:**
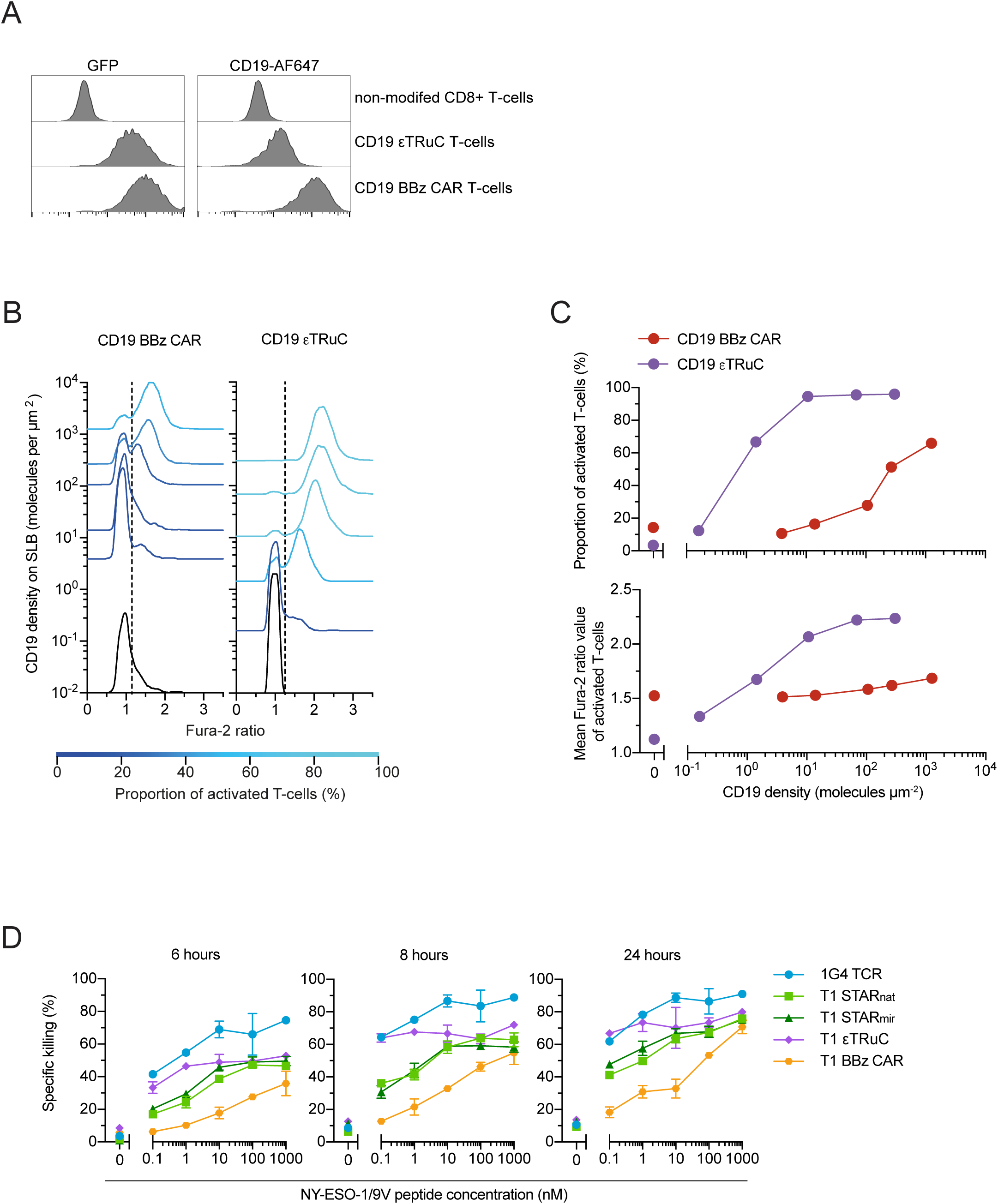
Calcium analysis of T-cells modified with CD19 BBz CAR- and CD19 εTRuC constructs. (**A**) Flow cytometric analysis of the CD19-antigen receptor surface expression on CD19 εTRuC- and CD19 BBz CAR T-cells. Translation levels of lentivirally encoded receptor constructs were correlated with the expression of GFP C-terminally linked via a T2A site (see Materials and Methods section). (**B**) Population-wide analysis of the calcium response of CD19 BBz CAR- and CD19 εTRuC T-cells confronted with SLBs functionalized with CD19 at indicated densities. Dashed lines indicate the Fura-2 ratio threshold above which T-cells were considered activated. (**C**) Upper panel: rendering of data shown in (B) as antigen dose response curves. Lower panel: mean Fura-2 ratio values (taken from B) measured for the activated fraction of CD19 BBz CAR- and CD19 εTRuC T-cells confronted with SLBs featuring CD19 at indicated densities. (**D**) *Ex vivo* cytotoxic capacity of antigen receptor-engineered T-cells. Indicated effector cells were cocultured at a ratio of 1:1 and for indicated times with K562-A2 luciferase-expressing feeder cells pre-pulsed with the NY-ESO-1 peptide derivative 9V at indicated concentrations. Error bars = sem from triplicates.

### TCCs but not second-generation CARs promote CTL-mediated target cell killing at low antigen levels

We next investigated the capacity of synthetic receptors to mediate target cell cytolysis in response to antigen. The use of the T1 system proved advantageous, as it allowed for gradual increase in antigen densities by titrating the NY-ESO-1 peptide to the target cell culture. As shown in **Fig. 4D**, we confronted T-cells, which had been modified with the 1G4 TCR, the T1 BBz CAR, T1 STAR_nat_, T1 STAR_mir_ or the T1 εTRuC as indicated, with 9V peptide-pulsed HLA-A2 expressing K562 cells at an effector to target ratio of 1:1. The degree of ensuing target cell lysis was measured after 6, 8 and 24 hours of co-culture.

Cytolysis was highly antigen-dependent in all cases. Overall, 1G4 TCR T-cells caused the highest degree of killing, especially at the lowest 9V peptide concentrations (0.1 nM) applied for pulsing (40, 65 and 65 percent of killing after 6, 8 and 24 hours, respectively). The killing response of T1 εTRuC-T-cells was equally sensitive (at 0.1 nM 9V peptide: 37, 63 and 67 percent of killing after 6, 8 and 24 hours, respectively), yet increased to a lesser extent at higher 9V peptide concentrations employed, even after 24 hours of co-culture. We consider it likely that the substantially higher affinity of T1-A2/9V may have interfered with synapse dissolution and - in its wake - serial killing.

At low antigen densities, killing responses of T-cells modified with the T1 STAR_nat_ (19, 32 and 42 percent of killing after 6, 8 and 24 hours, respectively) and the T1 STAR_mir_ (20, 29 and 44 percent of killing after 6, 8 and 24 hours, respectively) were indistinguishable, yet scored significantly lower when compared to those caused by 1G4 TCR- and T1 εTRuC T-cells. Target cell lysis increased gradually with increasing antigen levels and time of co-culture. Lysis saturated at about the level that was induced by T1 εTRuC T-cells at 9V peptide concentrations of 10 nM and higher. Our measurements imply that signaling in response to low levels of the high affinity A2/9V antigens was most robust for T-cells featuring the T1 εTRuC, which exceeded that of T-cells expressing other TCR/CD3-based receptor constructs.

In line with the calcium- and IFN-γ-based readouts of T-cell responsiveness, T1 BBZ CAR T-cells showed by far the least efficient cytolytic response when confronted with target cells pulsed with the lowest 9V peptide concentration (0.1 nM). Target cell killing was in fact indistinguishable from that of the negative control measured 6 and 8 hours after cell-pooling. After 24 hours of co-culture specific cytolysis increased to 6% after subtraction of background activity. Only at the highest antigen dose applied (1 µM) target cell killing approached levels conferred by T-cells engineered with the 1G4 TCR or TCR/CD3-based synthetic antigen receptor constructs.

Taken together, TCRs and TCR/CD3-based synthetic constructs outperform conventional second-generation CARs at the level of sensitized target cell killing, a finding that is consistent with the earlier observed differences in dose-responses as measured by calcium signaling and IFN-γ-secretion.

### Superior membrane proximal signal transmission by TCR/CD3-based synthetic constructs

T-cell antigen recognition is intimately tied to synaptic receptor-antigen binding events. As outlined above, these become transmitted across the plasma membrane by means of Lck-mediated ITAM-phosphorylation which is followed by ZAP70’s recruitment to and activation at phospho-ITAMs for further amplification of downstream signaling. In view of the considerable differences in antigen detection capacities we had observed above, we sought to juxtapose signaling activities immediately downstream of T1 STAR_nat_, T1 STAR_mir_, T1 εTRuC and the second-generation T1 BBz- and CD19 BBz CARs. We included in our analysis RA14 TCR T-cells confronted with A2/CMV serving as a benchmark for single molecule antigen sensitivity.

To assess membrane-proximal downstream signaling, we first quantitated ITAM- and ZAP70-phosphorylation levels in T-cells responding to SLB functionalized with titrated antigen densities. To this end, receptor-modified T-cells confronted with stimulatory SLBs were fixed and subjected to membrane permeabilization. Relative levels of CD3ζ ITAM#2- and ZAP70-phosphorylation were then determined by TIRF-microscopy with the use of fluorescence-tagged specific monoclonal antibodies (mAbs) (**Fig. 5A**, left panel). We targeted phosphorylated CD3ζ ITAM#2 since the resulting immune fluorescence gave rise to the best signal to noise ratio when compared to the fluorescence obtained via CD3ζ pITAM#1- and pITAM#3-specific mAbs^11^. Levels of activated ZAP70 were monitored with the use of a fluorescence-labeled mAb specific for phosphorylated ZAP70 (pY319) (**Fig. 5A**, right panel) (for further details please refer to the Methods section).

**Figure 5:**
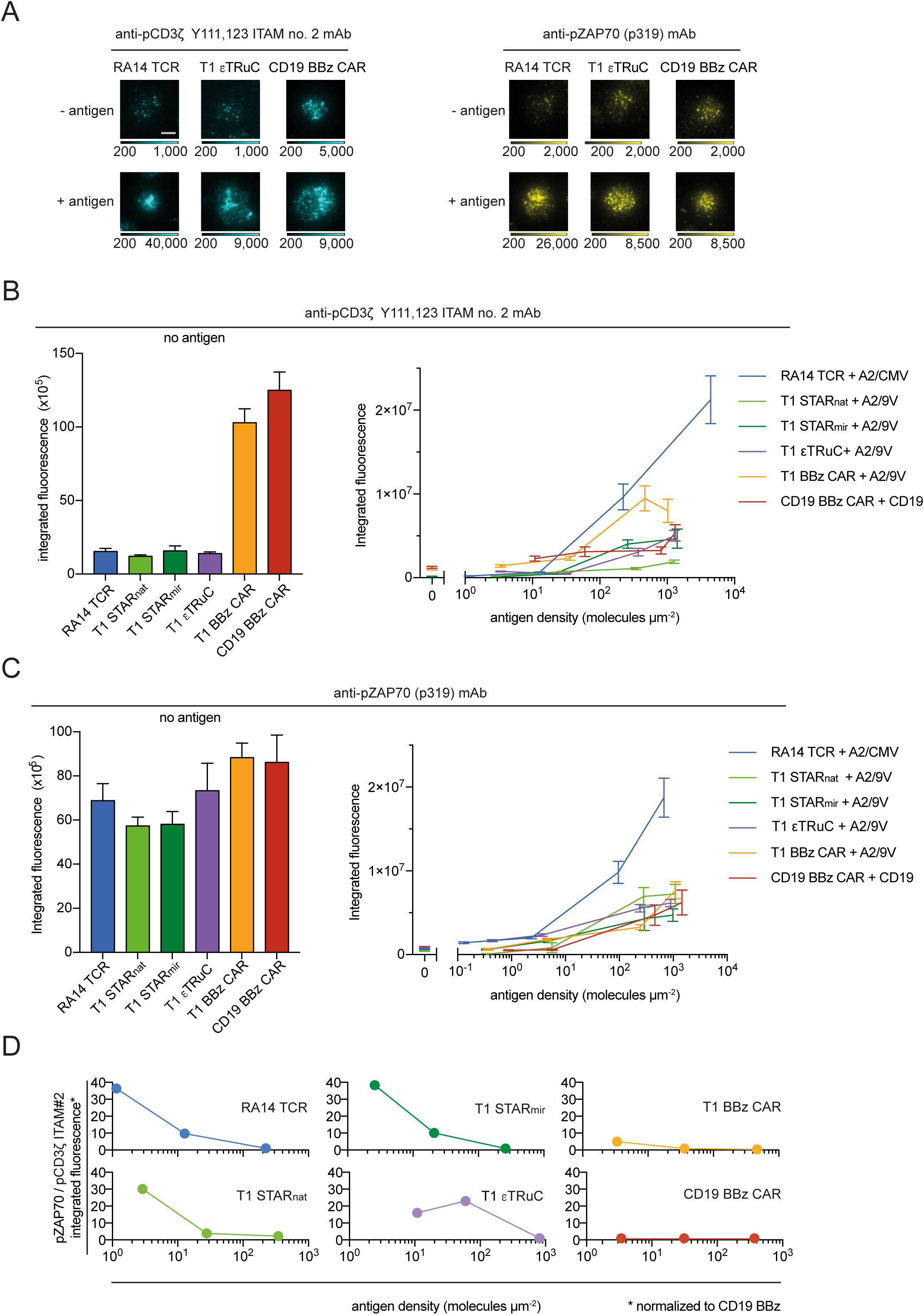
Immunofluorescence-based analysis of synapse-associated CD3ζ pITAM#2 and pZAP-70. (**A**) Synaptic immunofluorescence recorded in TIRF mode with the use of indicated antibodies. Shown are synapses of T-cells modified with indicated antigen receptors and in contact with SLBs featuring ICAM-1, B7-1 and, if indicated, their nominal antigen (A2/CMV, A2/9V and CD19, respectively). (**B**) Synapse-associated integrated fluorescence of CD3ζ Y111,123 ITAM no. 2 in the absence (left panel) and presence of antigen (right panel) as indicated. (**C**) Synapse-associated integrated fluorescence of ZAP-70 pY319 in the absence (left panel) and presence of antigen (right panel) as indicated. (**D**) Ratios of pZAP70 and pITAM#2 integrated fluorescence intensities which had been normalized to ratio values measured for CD19 BBz CAR T-cells. For analysis, integrated pZAP70 fluorescence values had been extrapolated for antigen densities associated with measured integrated pITAM#2 fluorescence values.

We noticed for both T1 BBz- and CD19 BBz-modified T-cells anti-CD3ζ pITAM#2 fluorescence signals in the absence of antigenic ligands, which amounted to levels that were up to ten times increased when compared to those obtained from all other T-cell lines analyzed (**Fig. 5B**, left panel). These observations are in good agreement with spurious calcium-signaling we had recorded in second-generation CAR T-cells but not in T-cells modified with TCRs or TCCs (**Fig 1H, I** , **Fig. 3** and **Ext. Fig. 5**).

Interestingly, for RA14 TCR-, T1 STAR_nat_-, T1 STAR_mir_- and T1 εTRuC T-cells we failed to detect a measurable increase in CD3ζ ITAM#2 -phosphorylation above background levels at antigen densities that were slightly above the thresholds for activation, i.e. sufficient to activate 60 to 90 % of the SLB-contacting T-cells (**Fig. 3, Ext. Fig. 6** and **Table 1**). Notably, we only observed a small increase in fluorescence signals for T-cells facing moderate antigen levels well above antigen thresholds. This implies that the number of triggered antigen receptors required to initiate full T-cell activation was significantly lower than the number of antigen receptors that were already partially phosphorylated in the absence of antigen. In contrast, at antigen levels well above thresholds for activation, we noticed a gradual rise in synaptic anti-CD3ζ pITAM#2 fluorescence for all T-cell lines analyzed. Yet in view of (i) the seemingly low number of triggered TCR/CD3-based constructs required to promote full T-cell activation and also because of (ii) the up to ten-fold differences in ectopic receptor expression levels (**Fig. 2B-E**), we reasoned that measured anti-CD3ζ pITAM#2 fluorescence intensities hardly constituted a reliable parameter for meaningful comparisons of antigen-receptor-proximal signaling output (see below). We considered them instead to be of analytic value only when viewed in the context of subsequent events in canonical TCR-signal transduction, in particular the recruitment and activation of ZAP70 to pITAMs of triggered antigen receptors.

Unlike the levels of synaptic CD3ζ pITAM#2 recorded in the absence of antigen, corresponding synapse-associated pZAP70 levels were only moderately higher in T1 or CD19 BBZ-CAR-T-cells than in RA14 TCR-, T1 STAR_nat_-, T1 STAR_mir_- and T1 εTRuC T-cells (**Fig. 5C**). We hence inferred that the seven to ten-fold higher CD3ζ pITAM#2 background levels associated with CAR-T-cells (**Fig. 5B**) resulted only in a 1.3 to 1.5-fold higher degree of ZAP70-activation. Such behavior would be consistent with a significantly weaker link between ITAM-phosphorylation and ZAP70-activation in second-generation CAR T-cells yet might also explain elevated calcium levels observed in the absence of antigen.

To gauge membrane-proximal amplification of downstream signaling in a fashion that was no longer compromised by varying levels of ectopic receptor expression, we determined the efficiency with which ZAP70 had become activated as a function of ITAM-phosphorylation. For this we divided synaptic anti-pZAP70 intensity values by anti-pITAM#2 fluorescence intensity values (**Fig. 5D**). Consistent with their extraordinary antigen sensitivity, RA14 TCR T-cells displayed the highest ratio, especially at low antigen densities, followed by T1 STAR_nat_, T1 STAR_mir_- and T1 εTRuC T-cells. Indicative of their limited detection capacities, CD19 and T1 BBz CAR T-cells exhibited by far the lowest pZAP70:pITAM#2 ratios at all antigen densities analyzed.

We therefore conclude that canonical progression from ITAM-phosphorylation to ZAP70-recruitment and -phosphorylation occurs at a much higher frequency in ultrasensitive RA14 TCR T-cells than in CD19 or T1 BBz CAR-T-cells, especially when antigen is present in low abundance. Of note, TCC-T-cells outperformed second-generation CAR-T-cells at all antigen densities tested.

## DISCUSSION

Empiric CAR designs elicit potent antitumor T-cell activity yet often fail to support long-term tumor remission due (i) to an inadequate sensitivity they convey towards TAAs and (ii) also the limited persistence of CAR-T-cell effector functions. Past efforts to address these shortcomings have rarely questioned the operational fitness of conventional CAR architectures but have instead hinged in large part on modifying or adding CAR modules^26–31^. Furthermore, receptor-ligand affinities, CAR stoichiometry and surface expression have been fine-tuned in previous attempts to boost overall performance.

Here we purposely abstained from reengineering one-dimensional CAR constructs. Our antigen receptor designs were instead guided by the rationale to emulate the unmatched fidelity and detection efficacy of conventional T-cells through the evolved TCR/CD3 architecture and its co-evolved signaling machinery. We reasoned that such hallmarks of antigen detection are at least in part conserved in TCR/CD3-based receptor systems as long as structural integrity is preserved. The observation that surface expression levels of all TCR/CD3-based receptor constructs scored significantly lower than those of endogenous or ectopically expressed TCR/CD3 complexes, underscores a need for further structural refinement. Nonetheless, the boosted functional performance of these constructs rivaled that of TCR/CD3 and hence corroborated our initial assumptions.

Our observations were consistent with previous reports^28, 29, 34, 35^, yet yielded in addition precise activation thresholds to be ranked with respect to the ultimate benchmark of a single antigen per synapse as set by CMV-specific RA14 TCR T-cells. Moreover, by focusing on A2/NY-ESO-1 as a molecular target to be recognized through (i) a TCR, (ii) a CAR or (iii) TCCs we have overcome complications arising from the otherwise divergent nature of epitopes recognized via TCR- and CAR T-cells and also sensitization of antigen recognition by co-receptor engagement. Furthermore, automated image recording and processing supported single cell-based analysis with single molecule resolution.

Notably, the T1 STAR_mir_ construct conferred high sensitivity towards ligands featuring a wide range of affinities. In contrast, both the T1 εTRuC and T1 STAR_nat_ failed to efficiently trigger T-cells in response to the low affinity A2/4D ligand with an off-rate falling within the range of most stimulatory TCR-pMHC bonds. We consider it likely that the T1 εTRuC’s rather flexible linker tethering the scF_v_ to CD3ε failed to efficiently transmit short-lived antigen-binding events. Initially, we suspected the low numbers of T-cell-resident T1 STAR_nat_ molecules to interfere with serial T1 STAR_nat_ engagement and receptor triggering by limiting A2/4D. However, since T-cell responses were not significantly affected by the ablation of endogenous TCR genes and the ensuing increase in STAR_nat_-surface expression, we consider it more likely that the observed inferior trigger rate resulted from deficits in receptor structure rather than limiting receptor surface expression. As mentioned earlier, the evolutionary conserved structure of the TCR/CD3 complex defines ligand engagement in an anisotropic fashion^33^, which is why we opted to design the T1 STAR_nat_ along with the T1 STAR_mir_ construct. However, the T1 STAR_nat_ design entailed pairing the V_H_- (V_L_) domain of the T1 scF_V_ with C_α_ (C_β_) and not with the evolutionary more related C_β_ (C_α_) -domain of the TCR, which may have negatively affected structural integrity. Minor structural shortcomings may indeed show the largest adverse impact in response to low affinity ligands. In contrast, the more conserved structure of T1 STAR_mir_ is consistent with the receptor’s high signaling capacity in response to A2/4D ligands and evolutionarily optimized transduction across the plasma membrane. As suggested by numerous studies, an orientation-selective mode of antigen engagement may reflect on specific geometric constraints governing the extraordinary TCR/CD3 signaling capacity ^22, 36, 37^, a view which is consistent with our finding that T1 STAR_nat_ T-cells scored higher than T1 STAR_mir_ T-cells in response to high affinity A2/9V and medium affinity A2/6T ligands (**Table 1**).

Unlike what has been described for the recognition of pMHC class complexes by most CD8+ T-cells, the sensitizing effect of CD8-coreceptor engagement proved in our hands only minor for T-cells modified with TCCs. However, none of the T-cell lines modified with TCCs approached the sensitivity benchmark set by RA14 TCR T-cells. Mounting evidence suggests that downstream signaling resulting from more stable TCR-antigen interactions with off-rates similar to those of T1-A2/9V and -A2/6T is less dependent on CD8-MHC class I binding ^38, 39^. We also consider it possible that CD8-mediated sensitization rests on the specific geometric framework determined by pMHC-engagement via CD8 and TCR/CD3, which may not be equally supported by T1 STAR_nat_, T1 STAR_mir_ and T1 εTRuC. Of note, mechanisms underlying αβTCR-mediated T-cell activation appear to serve in particular the detection of low affinity pMHC antigens present in low abundance and outnumbered by structurally similar yet non-stimulatory pMHCs. In contrast, the related γδTCRs, B-cell- and Fc-receptors act all in a coreceptor-independent fashion: unlike αβTCRs, they appear optimized for the recognition of more abundant ligands which feature higher affinities and demand in turn - to maintain a high fidelity of detection - a lesser degree of discrimination from other proteins ^40, 41^. The fact that TAA-recognition by TCC formats are almost insensitive to CD8-co-engagement, places them closer to γδTCRs. Most importantly, it qualifies them for their use in advanced anti-tumor therapies targeting low or heterogeneously expressed TAAs other than peptide-loaded MHC class I molecules.

Unlike CARs, TCR/CD3-based synthetic constructs did not give rise to antigen-independent signaling when expressed in T-cells. In principle, all domains of the CAR can contribute to spurious activation. Replacing the CD28-derived costimulatory domain with that of 4-1BB was reported to reduce tonic signaling ^11^, yet even 20% of T-cells modified with 4-1BB-based CARs displayed elevated intracellular calcium levels in the absence of antigen, a behavior we did not observe in T-cells engineered with TCCs. Another reason underlying antigen-independent signaling in CAR T-cells may involve cluster formation of CAR molecules induced by dimerization or oligomerization of CAR-associated scF_V_. Because of the lack of constant domains and their stabilizing inter-domain disulfide bonds, a diminished structural integrity of V_H_- and V_L_-domain may promote in many scF_V_s the formation of interchain V_H_-V_L_-pairing and - as a consequence - daisy-chained CAR entities. These may undergo spontaneous phosphorylation in the absence of any antigen-mediated trigger. Of note, all TCCs employed in our study featured a scF_V_ -based antigen-targeting domain as well, but did not allow for spurious signaling. Underlying reasons may involve the - compared to CARs - up to 34-times lower surface levels of TCCs, which may render antigen-independent receptor clustering and ensuing signaling less likely. Extensive ER-quality control during TCR/CD3 assembly ^19^ may be reflective of substantial evolutionary pressure to preclude spurious signaling and to allow in turn for sensitized detection of single antigens. In support of this notion is the observed recruitment of the inhibitory kinase Csk via an intracellular CD3ε-motif ^42^. When incorporated into the cytoplasmic tail of a 28zCAR, this short peptide sequence appeared to reduce spurious activation ^42^.

Large variations in receptor expression levels in addition to the considerable degree of antigen-independent CD3ζ ITAM background phosphorylation complicated in our study the immunofluorescence-based identification of membrane-proximal mechanisms which underlie the observed functional differences between TCRs, CARs and TCCs. Of note, at antigen levels that were close to activation thresholds, we failed to observe in any of the investigated receptor-engineered T-cells an increase in ζITAM-phosphorylation above background levels. From this we infer that antigen-induced phosphorylation of only few antigen receptors leads to the recruitment and activation of ZAP70 in numbers that are already sufficient for T-cell activation. For this to occur, the phosphorylation status of individual antigen-triggered TCR/CD3 complexes, TCCs and CARs likely exceed significantly that of otherwise highly abundant yet only partially phosphorylated receptor entities, as they may result in the absence of antigen from sporadic Lck-interactions.

Interestingly, in response to low antigen densities (ten or fewer antigens per square micron) synaptic ITAM-phosphorylation scored significantly higher in CAR T-cells than in RA14 TCR T-cells. However, only RA14 TCR-but not CAR-T-cells showed synaptic ZAP70-phosphorylation above background at these same antigen densities. Ratiometric analyses of anti-pITAM- and anti-pZAP70-immunofluorescence intensities imply that antigen-driven progression from ITAM phosphorylation to ZAP70 activation is markedly reduced in CAR T-cells. In the absence of direct evidence, we hypothesize that complete phosphorylation of all available ITAMs within a TCR/CD3 complex, a TCC or a CAR is critical for effective ZAP70-activation and further downstream signaling. This notion is supported by the observation that ZAP phosphorylation depends on stable ZAP70-association with the cytoplasmic tails of CD3, which relies in turn on the dynamic engagement of both ZAP70-SH2-domains with neighboring phospho-tyrosine residues ^43^. Since levels of activated ZAP70 are undetectable for T-cells undergoing already robust calcium signaling, we presume that only few catalytically active ZAP70 molecules are required for T-cell activation.

Our findings justify further *in vivo* pre-clinical testing of TCC-modified T-cells in murine *in vivo* cancer models. In fact, recent studies have suggested superior *in vivo* anti-tumor activity of CD19 εTRuC ^28^ - and CD19 STAR ^29, 35^ - modified T-cells. Yet without exact knowledge of target cell-expressed CD19 levels, the extent to which CD19 εTRuC- or CD19 STAR-modified T-cells achieve eradication of CD19^low^ tumor cells, as they may emerge under selective pressure following CAR-T-cell therapy, remains to be identified.

## MATERIALS AND METHODS

### Constructs for lentiviral transduction

HLA.A2/NY.ESO-1-specific lentiviral constructs were generated by Gibson assembly and cloned in lentiviral vector p526. Sequences from the T1 scF_V_, the c58c61 1G4 TCR and the CD19-Kymriah CAR construct ^6, 37, 44^ were cloned into the lentiviral vector epHIV7. T1 CAR BBz was generated by exchanging the FMC63 single-chain variable fragment (scF_V_) by the T1 scF_V_ within the CD19 CAR BBz construct. T1 εTRuC and CD19 εTRuC were generated by tethering the T1 scF_V_ and FMC62 scF_V_ to CD3ε, respectively, via the flexible linker (G_4_S)_3_. T1 STAR_nat_ was generated by directly tethering T1-V_H_ to the murine TCRα constant region (Uniprot: A0A075B662) and T1-V_L_ to the murine TCRβ constant region (Uniprot: A0A075B5J4). T1 STAR_mir_ was generated by directly tethering T1-V_L_ to the murine TCRα constant region and T1-V_H_ to the murine TCRβ constant region. All lentiviral constructs contained a copGFP transduction marker separated by a viral T2A cleavage sequence ^45^. Please refer to the appendix for DNA sequences encoding NY-ESO-1-specific synthetic receptors.

### Generation of lentiviral and CRISPR-Cas9 specificity-redirected T-cells

CD8^+^ T-cells were isolated from HLA.A2^-^ donors using magnetic-activated cell sorting (MACS, Miltenyi Biotec). Anti-CD3/CD28/CD2 Immunocult (Stemcell Technologies) was used for T-cell activation. T-cells were cultivated at 37°C and in 5% atmospheric CO_2_ in Roswell Park Memorial Institute (RPMI) 1640 medium supplemented with 25 mM HEPES, 10% human serum, 100 U ml^−1^ penicillin/streptomycin, 50 µM of 2-mercaptoethanol (Thermo Fisher Scientific) and 50 U ml^−1^ recombinant interleukin-2 (IL-2; Novartis).

For lentiviral transduction, polybrene (Sigma Aldrich) was added to the medium containing the lentivirus on the following day at a concentration of 8 µg ml^−1^ to stably integrate synthetic receptor-encoding genes. On day 7 post transduction the percentage of transduced cells was determined for each condition by assessing copGFP expression by flow cytometry. Transgene-positive T-cells were enriched using fluorescence activated cell sorting (FACS) (Sony SH800).

For CRISPR-Cas9-mediated replacement of the TCR, 300 U/ml IL-2 was added to the medium of the CD8^+^ T-cells on the day after Immunocult activation. TCRα-chain and TCRβ-chain were knocked-out as described^45^. In brief, *TRAC* and *TRBC* were targeted using ribonucleoprotein (RNP) based on the following CRISPR RNA (crRNA) (Integrated DNA Technologies): 5′- GGAGAATGACGAGTGGACCC-3′ for *TRBC* (targeting both *TRBC1* and *TRBC2*) and 5′- AGAGTCTCTCAGCTGGTACA-3′ for *TRAC*. crRNA was incubated with trans-activating RNA (tracrRNA) (Integrated DNA Technologies), both at 80 µM and at 95°C for 5 minutes, then cooled down to 25°C, after which Cas9 was added. Cas9-RNP mix was stored on ice prior to use. CRISPR- Cas9-mediated knock-in of the RA14 or 1G4 TCR was performed as described ^32^. Double-stranded DNA PCR products containing the TCRβ-chain, T2A, TCRα-chain, stop codon and a poly-A flanked by the *TRAC*-specific homology arms were produced via polymerase chain reaction (PCR). 5x10^6^ T- cell were electroporated (Amaxa Nucleofector II) in 100 µl electroporation buffer (Lonza) in the presence of Cas9-RNP and 2 µg of DNA-template. Cells were cultured in the presence of 180 U/ml IL-2. On day 7 of the procedure TCR replacement was confirmed using anti-RA14 or anti-1G4 tetramer staining. Genetically modified T-cells were enriched via FACS (Sony SH800).

For expansion, HLA.A2/NY-ESO-1-specific cells were cocultured with K562/HLA.A2 cells which had been irradiated with 80 Gy and pulsed with 100 nM NY.ESO-1/9V peptide. RA14 TCR T-cells were cocultured with K562/HLA.A2 cells irradiated with 80 Gy and pulsed with CMV peptide (NLVPMVATV), and CD19-specific cells were cocultured with K562/CD19 cells added at a ratio of 4:1 and irradiated with 80 Gy. Every second day, half of the medium was exchanged with fresh T-cell medium containing recombinant IL-2 (50 U ml^−1^). Experiments were conducted at day 7-10 of the expansion protocol and 16 or more hours after medium exchange.

For generation of T1 STAR_nat_ and T1 STAR_mir_ TCRαβ^-^ cells, T-cells were first subjected to lentiviral transduction and one round of expansion as described above. Next, T-cells were re-stimulated with Immunocult in preparation for CRISPR-Cas9-mediated TCR-knock-out as described above. For generation of RA14 TCR/T1 BBz CAR and RA14/ T1 εTRuC double positive T-cells, T-cells were either first subjected to CRISPR-Cas9-mediated TCR-exchange and then expanded as described above, or first subjected to lentiviral transduction and then to CRISPR-Cas9-mediated TCR-exchange. T-cells were then re-stimulated with Immunocult for lentiviral transduction as described above.

### Quantitation of cell-surface synthetic receptor molecules

Engineered T-cells were stained with increasing concentrations of AF647-conjugated anti-human TCR α/β antibody (IP26: Biolegend), HLA.A2/9V-AF647 or CD19-AF647 to determine the concentration of the staining agent leading to label saturation for CMV-specific RA14 TCR T-cells, HLA.A2/NY-ESO-1-specific engineered T-cells and CD19-specific CAR T-cells. The mean fluorescence intensity (MFI) of each cell population was measured by flow cytometry. The MFI of AF647-conjugated quantitation beads (Quantum MESF; Bangs laboratories) was determined to give rise to a calibration curve allowing to standardize fluorescence intensity (molecules of equivalent soluble fluorochrome, MESF). MFI values of cell populations were converted to MESF units according to manufacturer’s instructions.

### Protein expression and purification

Codon-optimized cDNAs encoding the extracellular domains excluding the leader sequence of HLA-A2 (UniProt: P01892) and beta-2-microglobulin (UniProt: P61769) were cloned into pET-28b and pHN1, respectively. A DNA sequence coding for a polyhistidine tag, which harbored 12 consecutive histidine residues (H_12_), was introduced C-terminally to HLA.A2 within pET28b. HLA.A2-H_12_ and beta-2-microglobulin were expressed in *Escherichia coli* BL21(DE3) as inclusion bodies and refolded in the presence of peptide to give rise to HLA.A2-peptide complexes ^46, 47^. After completion of the refolding reaction, 300 ml of protein solution was dialyzed three times against 10 l PBS. Correctly conformed HLA.A2-peptide complexes were purified using Ni^2+^-NTA agarose chromatography (HisTrap excel, 2 × 5 ml^2^ columns connected in series; GE Healthcare Life Sciences) followed by size-exclusion chromatography (Superdex 200 10/300 GL; GE Healthcare Bio-Sciences). cDNAs encoding the extracellular portions of human proteins ICAM-1 (UniProt: P05362) and B7-1 (UniProt: P33681) without the leader sequence was PCR-amplified and cloned into the pAcGP67 of the BaculoGold Baculovirus Expression System (BD Biosciences). This vector was modified so that all proteins featured a C-terminal H_12_-tag. We produced virus using the Baculovirus Expression system according to the manufacturer’s instructions. High 5 cells (BTI-Tn-5B1-4; Thermo Fisher Scientific) were infected with virus particles. The supernatant was collected 4 days after infection at 90% cell viability. After centrifugation and filtration (0.45 µm) it was dialyzed against PBS by tangential flow filtration (Minimate Tangential Flow Filtration System equipped with a 10 kDa T-Series cassettes; Pall Corporation). The supernatant containing the proteins was concentrated eightfold and then diluted eightfold with PBS for a total of three times for buffer exchange. The processed supernatant was concentrated once more eightfold and then subjected to Ni^2+^-NTA agarose chromatography. Protein was eluted with PBS/300 mM imidazole (pH 7.4) and subjected to size-exclusion chromatography (Superdex 200 increase 10/300 gl and Superdex 75 Increase 10/300 GL, GE Healthcare Life Sciences) and anion-exchange chromatography (Mono Q 5/50 GL; GE Healthcare Life Sciences). The extracellular portion of CD19 (UniProt: P15391) was C-terminally fused to a polyhistidine tag containing 10 hystidine residues (H_10_), expressed in HEK cells and purified via Ni^2+^-NTA agarose affinity chromatography followed by size-exclusion chromatography and anion-exchange chromatography ^48^.

All purification steps were performed on an ÄKTA pure chromatography system (GE Healthcare Life Sciences). For all proteins, the fractions containing properly conformed proteins were identified using SDS-polyacrylamide gel electrophoresis (PAGE) followed by silver staining. Immediately after purification, protein was labeled and/or adjusted to 50% glycerol/PBS or snap-frozen in liquid N_2_ for storage at -80°C.

### Fluorophore conjugation

200 μg of HLA.A2-peptide complex in PBS was concentrated to 1 mg ml^−1^ with the use of Amicon Ultra Centrifugal Filters (Merck), adjusted to pH 8.3 by adding freshly prepared NaHCO_3_ (0.1 M final concentration) and incubated first for 30 minutes at room temperature, which was followed by an overnight incubation at 4°C with *N*-hydroxysuccinimide ester derivatives of AF647 (Thermo Fisher Scientific) present in two-fold molar excess.

### SLB preparation

Supported lipid bilayers (SLBs) were prepared as described ^49^. In brief, 1,2-dioleoyl-*sn*-glycero-3-[N(5-amino-1-carboxypentyl)iminodiacetic acid)succinyl] (DGS-NTA(Ni)) and 1-palmitoyl-2-oleoyl-*sn*-glycero-3-phosphocholine (POPC) (both from Avanti Polar Lipids) were dissolved in chloroform and mixed at a 1:50 molar ratio. Lipids were dried under vacuum overnight in a desiccator and then resuspended in 10 ml of degassed PBS and sonicated in a water bath sonicator (Q700; Qsonica) under nitrogen at 120-170 W for at least 60 minutes until the suspension had lost turbidity. To remove non-unilamellar vesicles, the lipid suspension was centrifuged for 1 hour at 37,000 rpm (150,000 g) at room temperature using a Sorvall RC M150GX ultracentrifuge with a S150AT-0121 rotor (Thermo Fisher Scientific). The supernatant was obtained and centrifuged again for 8 hours at 43,000 rpm (288,000 g) at 4°C employing the same rotor and centrifuge. The supernatant was filtered (0.2 µm) (Filtropur S 0.2; Sarstedt) and stored at 4°C for up to 6 months.

Glass slides (22 x 64 mm^2^ no. 1.5 borosilicate; Menzel-Gläser) were plasma cleaned (Zepto, Diener Electronic) for 15 minutes and attached with the use of dental imprint silicon putty (Picodent twinsel 22, Picodent) to a Lab-Tek 12-well chamber (ThermoFisher Scientific) from which the original glass bottom had been removed. Slide-exchanged chambers were then incubated with lipid vesicle suspension for at least 10 minutes, after which they were extensively rinsed with PBS. For functionalization, SLBs were incubated for 60 minutes with protein-containing PBS and then rinsed twice with 15 ml PBS to remove unbound protein.

### Surface plasmon resonance (SPR) for T1 scFv affinity and kinetics

T1 scFv were produced by Absolute Antibody Ltd.. SPR experiments were performed with a Biacore T200 instrument (GE Healthcare). All experiments were conducted in degassed and filtered HBS-EP, at 37°C. Chip activation was performed using following parameters:

| Compound | Sample volume | Flow rate | Contact time |
| --- | --- | --- | --- |
| EDC / NHS | 200 μl | 10 μl min<sup>-1</sup> | 600 s |
| 10 μL Streptavidin in 190 μL Acetate (pH 4.5) | 200 μl | 10 μl min<sup>-1</sup> | 600 s |
| ProteON | 200 μl | 10 μl min<sup>-1</sup> | 300 s |
| Glycine buffer (pH 2.5) | 200 μl | 10 μl min<sup>-1</sup> | 300 s |

Refolded and biotinylated pMHCs were immobilized on a CM5 S series sensor chip at a flow rate of 10 µl min^-1^. Surface densities were set to roughly 300 or 1000 resonance units (RU) for kinetic and affinity measurements, respectively. Recombinant CD58 (biotinylated) was used as a negative control in the reference cell at a similar surface density, due to its comparable size. Flow cells were blocked by injecting d-Biotin twice at a concentration of 250 µM for 60 seconds at a flow rate of 10 µl min^-1^. The system was equilibrated by doing 10 start-up cycles that resembled the parameters for flow rate and contact time for the consecutive sample measurements. Recombinant scF_V_s were injected into the system using the “low consumption rate” at a flow rate of either 1 µl s^-1^ or 10 µl s^-1^ for either 20 seconds or 180 seconds. Ten different dilutions were injected consecutively into the system, followed by a dissociation step that lasted for 300 seconds.

For the determination of binding affinities, RUs were measured at the end of the association phase of a given concentration of the recombinant protein. RUs were plotted against the concentration of the recombinant protein and curve fitting was done with GraphPad Prism 7.04 with the “One Site – specific binding” function. For the determining of dissociation rates, a “One Phase Decay” function was fit to the data.

### Microscopy setup

Microscopy was conducted using two inverted setups. One setup allowed for TIR-based imaging and was built around an Eclipse Ti-E microscope body (Nikon Instruments) that was equipped with a chromatically corrected 100× TIR objective (CFI SR Apo TIR 100× Oil NA:1.49; Nikon Instruments), a 647 nm diode laser (OBIS) for excitation and a custom-made Notch filter (Chroma Technology) to block reflected stray light of 647 nM from reaching the camera. The setup was furthermore equipped with an ET700/75 emission bandpass filter (Chroma Technology) present in the emission pathway and a dichroic (QUAD Cube). An iXon Ultra 897 EMCCD camera (Oxford Instruments) was used for data recording. An 8-channel DAQ-board PCI-DDA08/16 (National Instruments) in combination with the microscopy automation and image analysis software MetaMorph (v. 7.8.13.0) (Molecular Devices) were used to program and apply timing protocols and control all hardware components of the microscope components.

A second inverted microscope (DMI4000; Leica Microsystems) was equipped with a 20× objective (HC PL FLUOTAR 20×/0.50 PH2∞/0.17/D, Leica Microsystems) and with a mercury lamp (EL6000; Leica Microsystems) for Fura-2-based calcium recordings. This microscope was equipped with a fast filter wheel containing 340/26 and 387/11 excitation bandpass filters (both Leica Microsystems). Data were recorded using a sCMOS Andor Prime95b (Photometrix). Open-source software µManager was used to program and control all hardware components.

### Measurements of antigen densities on SLBs

SLB antigen density was determined by counting the number of diffraction-limited fluorescent events within a region of interest (ROI) or by dividing the fluorescence intensity value within a ROI by the single-molecule fluorescent intensity value. For bilayers featuring clearly distinguishable diffraction-limited fluorescence events, 30 images were recorded within a ROI of 100 x 100 pixels. The total number of molecules within the ROI of each image was determined using the Fiji Thunderstorm plugin (ImageJ/Fiji) and corrected for pixel size and number of images to determine the antigen density (1 pixel ≙ 0.0256 µm^2^, 100 × 100 pixels = 10,000 pixels ≙ 256 µm^2^). For determining antigen densities of SLBs the average single-molecule fluorescence intensity value of at least 300 molecules was determined within the ROI using the Fiji Thunderstorm plugin as described above. The average integrated intensity value of ROIs of 10 images was determined and divided by the average single-molecule intensity value to arrive at the number of molecules within a given ROI. Finally, this value was corrected for pixel size to determine antigen density. To quantitate low antigen densities giving rise to distinct diffraction-limited fluorescence events, we counted the number of events within a given ROI for at least 30 images and divided the average number by the area of the ROI. Fluorophore densities were lastly converted to antigen densities by factoring in the protein-to-dye ratio determined by spectrophotometry.

### Determination of mobile fraction using FRAP

For determination of the SLB-resident mobile protein fraction we employed fluorescence recovery after photo bleach (FRAP) as described ^49^. A circular aperture was placed in the beam path. We recorded three images prior to bleaching the defined area of the bilayer and another ten images after the bleach pulse at 1-minute time intervals. Fluorescence intensity values recorded after bleaching were normalized against the average of the fluorescence intensity values recorded prior to photobleaching.

### Calcium imaging

Intracellular changes in Ca^2+^ levels were measured using the ratiometric calcium-sensitive dye Fura-2-AM as described ^50^. A total of 9x10^5^ cells were incubated in 0.5 ml imaging buffer (HBSS, supplemented with 2mM CaCl2, 2mM MgCl2, 2% FCS) supplemented with 5 µM Fura-2-AM for 15 minutes at 37°C, washed twice with 10 ml of imaging buffer and resuspended in 135 µl imaging buffer. Cells were kept at room temperature for a maximum of 30 minutes prior to starting the experiment. Immediately before imaging, PBS was exchanged for imaging buffer. T-cells were pipetted into the imaging buffer and allowed to settle for 30 seconds after which 510/80 nm emission was recorded while rapidly switching between 340 nm and 387 nm excitation for 20 minutes at time intervals of 15 seconds.

An inhouse custom-built Matlab software was used to track cells in each frame using a published particle tracking^23^. Tracking parameters were chosen so only single cells in contact with the SLB were included. We used the Matlab software to create ratio images for each frame. ‘Methods for automated and accurate analysis of cell signals’ or MAACS was used for population analysis as described^51^. For each trajectory within a population the ratio was normalized frame-wise to that of the population median of T-cells in contact with antigen-free SLBs. Cells that were above the threshold for at least 80% of their trajectory were plotted in a dose-response curve. The calcium histograms were compiled from the measured population values of the median Fura-2-AM ratio corresponding to the first 10 frames after the peak Fura-2-AM ratio value within the trajectory. The latter was normalized frame-wise to the population median of the negative control, i.e., cells confronted with antigen-free SLBs.

### Quantitation of cell-surface synthetic receptor molecule density

Engineered T-cells were stained with increasing concentrations of AF647-conjugated anti-human αβTCR antibody (IP26: Biolegend) and HLA.A2/9V-AF647 for determination of label saturation concentration of CMV-specific RA14 TCR T-cells and HLA.A2/NY-ESO-1-specific engineered T-cells, respectively. Labeled cells were dropped in imaging buffer on SLBs functionalized with ICAM-1 and allowed to settle for 8 minutes. Hereafter, images were recorded for at least 30 cells per condition. The average integrated intensity values of cells were determined, divided by the single-molecule intensity and corrected for pixel size and number of images to determine the antigen density.

### Quantitation of synapse-associated CD3**ζ** pITAM#2 and pZAP70

Pre-warmed T-cells were seeded in imaging buffer on SLBs and allowed to settle for 12 minutes at 37°C. T-cells were fixed for 30 minutes at room temperature in PBS supplemented with 20 mM NaF, 4% formaldehyde, 0.2% Triton X-100 Surfact-Amps (Thermo Fisher Scientific) and 2 mM Na_3_O_4_V. Formaldehyde was then gently washed away with washing buffer consisting of PBS supplemented with 20 mM NaF, 2 mM Na_3_O_4_V, 0.2% Triton X-100 and 3% bovine serum albumin (BSA). After three washes SLBs were incubated for 20 minutes at room temperature to block nonspecific binding. T-cells were incubated overnight at 4 °C with either mAb reactive to mouse phospho-CD247 (CD3 zeta) (Tyr111, Tyr123) (clone EM-55; Thermo Fisher Scientific) or mAb reactive to mouse ZAP70 phosphorylated at position 319 (clone 1503310; BioLegend). The next day, unbound antibody was washed away with washing buffer, incubated with goat anti-mouse IgG (H+L) AF647 (catalog no. A-21235; Thermo Fisher Scientific) and incubated for 3 hours at room temperature. Unbound antibody was washed away using washing buffer. Cells were stored at 4°C in the dark until acquisition of images. Integrated intensity values were recorded as described earlier with the exception that cells had been fixed.

### Quantitation of IFN-**γ** secretion after antigenic stimulation of T-cells on SLBs

IFN-γ production was quantitated through enzyme-linked immunosorbent assay (ELISA) of supernatants of 30,000 T-cells that had been stimulated through antigen-functionalized SLBs harboring different ligand densities. Prior to stimulation, SLB antigen density was determined after which buffer was exchanged to RPMI 1640 medium supplemented with 25 mM of HEPES, 10% FCS, of penicillin/streptomycin (100 U ml^-1^), 2 mM of L-glutamine and 50 µM of 2-mercaptoethanol. Hereafter, T-cells were seeded and incubated for 72 hours. The amount of secreted IFN-γ was determined the LEGEND MAX™ Human IFN-γ ELISA Kit (BioLegend) according to the manufacturer’s instructions.

### Killing assay of peptide-pulsed K562

HLA.A2^+^, luciferase firefly-BFP^+^ K562 cells were pulsed with different concentrations of HLA.A2/9V at 37°C for 1 hour. Cells were washed and resuspended in RPMI 1640 medium supplemented with 25 mM HEPES, 10% human serum, 100 U ml^−1^ penicillin/streptomycin, 50 µM of 2-mercaptoethanol (Thermo Fisher Scientific) and 75 µg ml^−1^ D-luciferin firefly (Biosynth). Cytotoxicity by engineered T-cells was determined after co-culturing peptide-pulsed K562 cells at an effector to target ratio of 1:1. Cells were co-cultured at 37 °C under 5% atmospheric CO_2_. K562 cultured in 40% DMSO were used as a 100% killing control. Relative light units indicating K562 viability were measured after 6, 8 and 24 hours on the microplate reader (Mithras LB 940). Specific cell lysis was determined as follows:

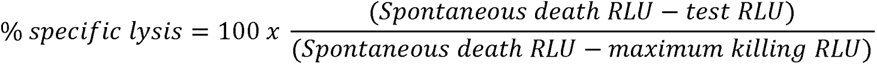

## Supporting information

https://drive.google.com/drive/folders/1J0S95Sg1-U1Gx6-xPFKRo2qXDpwHySQ7

## ACKNOWLEDGEMENTS

T.P. and J.B.H. received funding from. V.M., J.G., R.P. and A.P. were supported by the Austrian Science Fund (FWF, grant XXX, grant XXX, grant XXX, grant XXX).

## AUTHOR CONTRIBUTIONS

T.P. and J.B.H conceived the project and wrote the ms.. T.P. conducted most experiments. V.P. and A.P. conducted critical experiments. R.M.H.V.C., W.W.S. and S.M. designed and provided antigen receptor constructs as well as T-cell lines. R.P. and J.G. contributed important expertise and software code. I.D.P. implemented microscopy hardware and provided important ideas. B.S. and J.S.F. provided kinetic data for T1-ligand interactions. V.D.R.G, M.H., M.F., O.D., K.S., D.H.B. and H.S. provided important insights.

## REFERENCES

1. Brentjens, R.J. et al. CD19-targeted T cells rapidly induce molecular remissions in adults with chemotherapy-refractory acute lymphoblastic leukemia. Sci Transl Med 5, 177ra138 (2013).

2. Fry, T.J. et al. CD22-targeted CAR T cells induce remission in B-ALL that is naive or resistant to CD19-targeted CAR immunotherapy. Nat Med 24, 20–28 (2018).

3. Kochenderfer, J.N., Yu, Z., Frasheri, D., Restifo, N.P. & Rosenberg, S.A. Adoptive transfer of syngeneic T cells transduced with a chimeric antigen receptor that recognizes murine CD19 can eradicate lymphoma and normal B cells. Blood 116, 3875–3886 (2010).

4. Maude, S.L. et al. Tisagenlecleucel in Children and Young Adults with B-Cell Lymphoblastic Leukemia. N Engl J Med 378, 439–448 (2018).

5. Porter, D.L. et al. Chimeric antigen receptor T cells persist and induce sustained remissions in relapsed refractory chronic lymphocytic leukemia. Sci Transl Med 7, 303ra139 (2015).

6. Maude, S.L. et al. Chimeric antigen receptor T cells for sustained remissions in leukemia. N Engl J Med 371, 1507–1517 (2014).

7. Turtle, C.J. et al. Immunotherapy of non-Hodgkin’s lymphoma with a defined ratio of CD8+ and CD4+ CD19-specific chimeric antigen receptor-modified T cells. Sci Transl Med 8, 355ra116 (2016).

8. Brudno, J.N. et al. T Cells Genetically Modified to Express an Anti-B-Cell Maturation Antigen Chimeric Antigen Receptor Cause Remissions of Poor-Prognosis Relapsed Multiple Myeloma. J Clin Oncol 36, 2267–2280 (2018).

9. Libert, D. et al. Serial evaluation of CD19 surface expression in pediatric B-cell malignancies following CD19-targeted therapy. Leukemia 34, 3064–3069 (2020).

10. Shah, N.N. & Fry, T.J. Mechanisms of resistance to CAR T cell therapy. Nat Rev Clin Oncol 16, 372–385 (2019).

11. Long, A.H. et al. 4-1BB costimulation ameliorates T cell exhaustion induced by tonic signaling of chimeric antigen receptors. Nat Med 21, 581–590 (2015).

12. Lynn, R.C. et al. c-Jun overexpression in CAR T cells induces exhaustion resistance. Nature 576, 293–300 (2019).

13. Gudipati, V. et al. Inefficient CAR-proximal signaling blunts antigen sensitivity. Nature Immunology 21, 848–856 (2020).

14. Huang, J. et al. A single peptide-major histocompatibility complex ligand triggers digital cytokine secretion in CD4(+) T cells. Immunity 39, 846–857 (2013).

15. Irvine, D.J., Purbhoo, M.A., Krogsgaard, M. & Davis, M.M. Direct observation of ligand recognition by T cells. Nature 419, 845–849 (2002).

16. Purbhoo, M.A., Irvine, D.J., Huppa, J.B. & Davis, M.M. T cell killing does not require the formation of a stable mature immunological synapse. Nat Immunol 5, 524–530 (2004).

17. Alcover, A., Alarcon, B. & Di Bartolo, V. Cell Biology of T Cell Receptor Expression and Regulation. Annu Rev Immunol 36, 103–125 (2018).

18. Dong, D. et al. Structural basis of assembly of the human T cell receptor-CD3 complex. Nature 573, 546–552 (2019).

19. Huppa, J.B. & Ploegh, H.L. In vitro translation and assembly of a complete T cell receptor-CD3 complex. J Exp Med 186, 393–403 (1997).

20. Call, M.E., Pyrdol, J., Wiedmann, M. & Wucherpfennig, K.W. The organizing principle in the formation of the T cell receptor-CD3 complex. Cell 111, 967–979 (2002).

21. Shah, K., Al-Haidari, A., Sun, J. & Kazi, J.U. T cell receptor (TCR) signaling in health and disease. Signal Transduct Target Ther 6, 412 (2021).

22. Zareie, P. et al. Canonical T cell receptor docking on peptide-MHC is essential for T cell signaling. Science 372 (2021).

23. Lo, W.L. et al. Lck promotes Zap70-dependent LAT phosphorylation by bridging Zap70 to LAT. Nat Immunol 19, 733–741 (2018).

24. Andreotti, A.H., Schwartzberg, P.L., Joseph, R.E. & Berg, L.J. T-cell signaling regulated by the Tec family kinase, Itk. Cold Spring Harb Perspect Biol 2, a002287 (2010).

25. Li, W. et al. Chimeric Antigen Receptor Designed to Prevent Ubiquitination and Downregulation Showed Durable Antitumor Efficacy. Immunity 53, 456–470 e456 (2020).

26. Helsen, C.W. et al. The chimeric TAC receptor co-opts the T cell receptor yielding robust anti-tumor activity without toxicity. Nat Commun 9, 3049 (2018).

27. Xu, Y. et al. A novel antibody-TCR (AbTCR) platform combines Fab-based antigen recognition with gamma/delta-TCR signaling to facilitate T-cell cytotoxicity with low cytokine release. Cell Discov 4, 62 (2018).

28. Baeuerle, P.A. et al. Synthetic TRuC receptors engaging the complete T cell receptor for potent anti-tumor response. Nat Commun 10, 2087 (2019).

29. Liu, Y. et al. Chimeric STAR receptors using TCR machinery mediate robust responses against solid tumors. Sci Transl Med 13 (2021).

30. Loffler, A. et al. Efficient elimination of chronic lymphocytic leukaemia B cells by autologous T cells with a bispecific anti-CD19/anti-CD3 single-chain antibody construct. Leukemia 17, 900–909 (2003).

31. Huppa, J.B. et al. TCR-peptide-MHC interactions in situ show accelerated kinetics and increased affinity. Nature 463, 963–967 (2010).

32. Schober, K. et al. Orthotopic replacement of T-cell receptor alpha- and beta-chains with preservation of near-physiological T-cell function. Nat Biomed Eng 3, 974–984 (2019).

33. Stewart-Jones, G. et al. Rational development of high-affinity T-cell receptor-like antibodies. Proc Natl Acad Sci U S A 106, 5784–5788 (2009).

34. Heitzeneder, S. et al. GPC2-CAR T cells tuned for low antigen density mediate potent activity against neuroblastoma without toxicity. Cancer Cell 40, 53–69 e59 (2022).

35. Mansilla-Soto, J. et al. HLA-independent T cell receptors for targeting tumors with low antigen density. Nat Med 28, 345–352 (2022).

36. Birnbaum, M.E. et al. Molecular architecture of the alphabeta T cell receptor-CD3 complex. Proceedings of the National Academy of Sciences of the United States of America 111, 17576–17581 (2014).

37. Adams, J.J. et al. T cell receptor signaling is limited by docking geometry to peptide-major histocompatibility complex. Immunity 35, 681–693 (2011).

38. Laugel, B. et al. Different T cell receptor affinity thresholds and CD8 coreceptor dependence govern cytotoxic T lymphocyte activation and tetramer binding properties. J Biol Chem 282, 23799–23810 (2007).

39. Liu, B. et al. 2D TCR-pMHC-CD8 kinetics determines T-cell responses in a self-antigen-specific TCR system. Eur J Immunol 44, 239–250 (2014).

40. Mallis, R.J. et al. Molecular design of the gammadeltaT cell receptor ectodomain encodes biologically fit ligand recognition in the absence of mechanosensing. Proc Natl Acad Sci U S A 118 (2021).

41. Deseke, M. & Prinz, I. Ligand recognition by the gammadelta TCR and discrimination between homeostasis and stress conditions. Cell Mol Immunol 17, 914–924 (2020).

42. Wu, W. et al. Multiple Signaling Roles of CD3epsilon and Its Application in CAR-T Cell Therapy. Cell 182, 855–871 e823 (2020).

43. Goyette, J. et al. Dephosphorylation accelerates the dissociation of ZAP70 from the T cell receptor. Proc Natl Acad Sci U S A 119 (2022).

44. Gao, Y. & Kilfoil, M.L. Accurate detection and complete tracking of large populations of features in three dimensions. Opt Express 17, 4685–4704 (2009).

45. Shagin, D.A. et al. GFP-like proteins as ubiquitous metazoan superfamily: evolution of functional features and structural complexity. Mol Biol Evol 21, 841–850 (2004).

46. Garboczi, D.N., Hung, D.T. & Wiley, D.C. HLA-A2-peptide complexes: refolding and crystallization of molecules expressed in Escherichia coli and complexed with single antigenic peptides. Proc Natl Acad Sci U S A 89, 3429–3433 (1992).

47. Clements, C.S. et al. The production, purification and crystallization of a soluble heterodimeric form of a highly selected T-cell receptor in its unliganded and liganded state. Acta Crystallogr D Biol Crystallogr 58, 2131–2134 (2002).

48. Lobner, E. et al. Getting CD19 Into Shape: Expression of Natively Folded "Difficult-to-Express" CD19 for Staining and Stimulation of CAR-T Cells. Front Bioeng Biotechnol 8, 49 (2020).

49. Axmann, M., Schutz, G.J. & Huppa, J.B. Single Molecule Fluorescence Microscopy on Planar Supported Bilayers. J Vis Exp, e53158 (2015).

50. Roe, M.W., Lemasters, J.J. & Herman, B. Assessment of Fura-2 for measurements of cytosolic free calcium. Cell Calcium 11, 63–73 (1990).

