## Supplementary material for "TCR/CD3-based synthetic antigen receptors (TCC) convey superior antigen sensitivity combined with high fidelity of activation": https://drive.google.com/drive/folders/1J0S95Sg1-U1Gx6-xPFKRo2qXDpwHySQ7

### Extended Data Figure 1

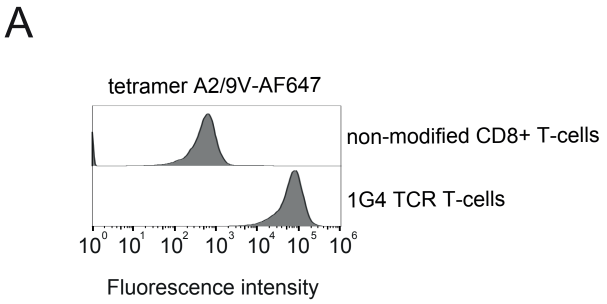


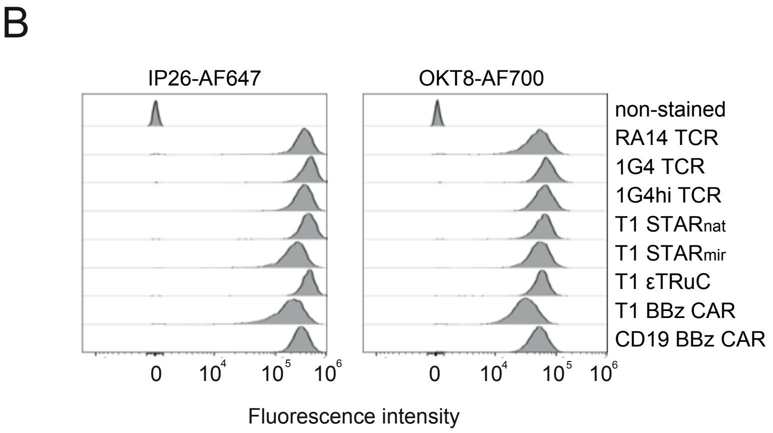


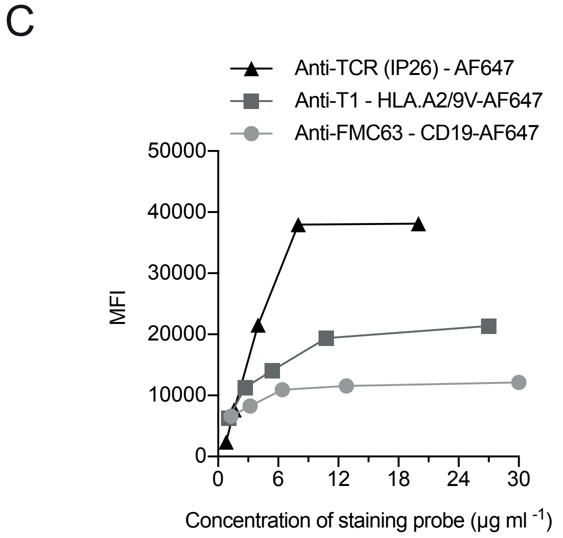


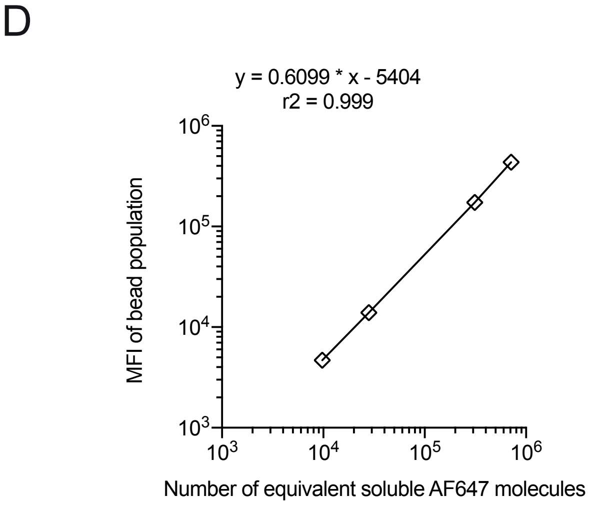

### Extended Data Figure 2

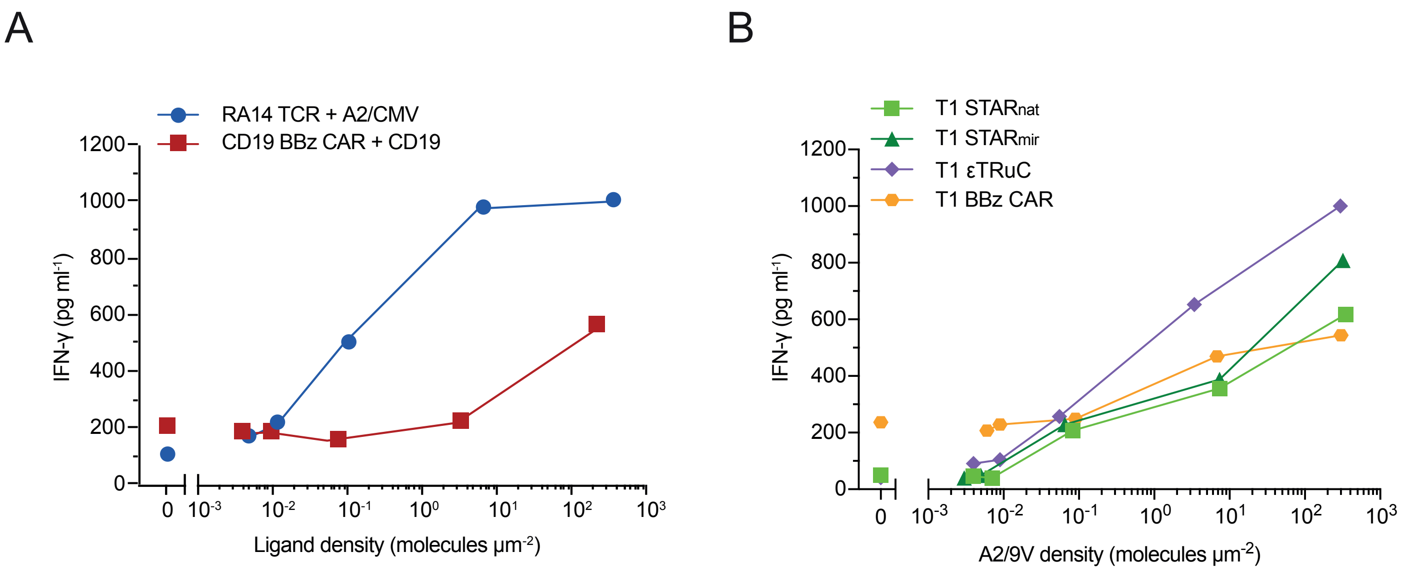

### Extended Data Figure 3

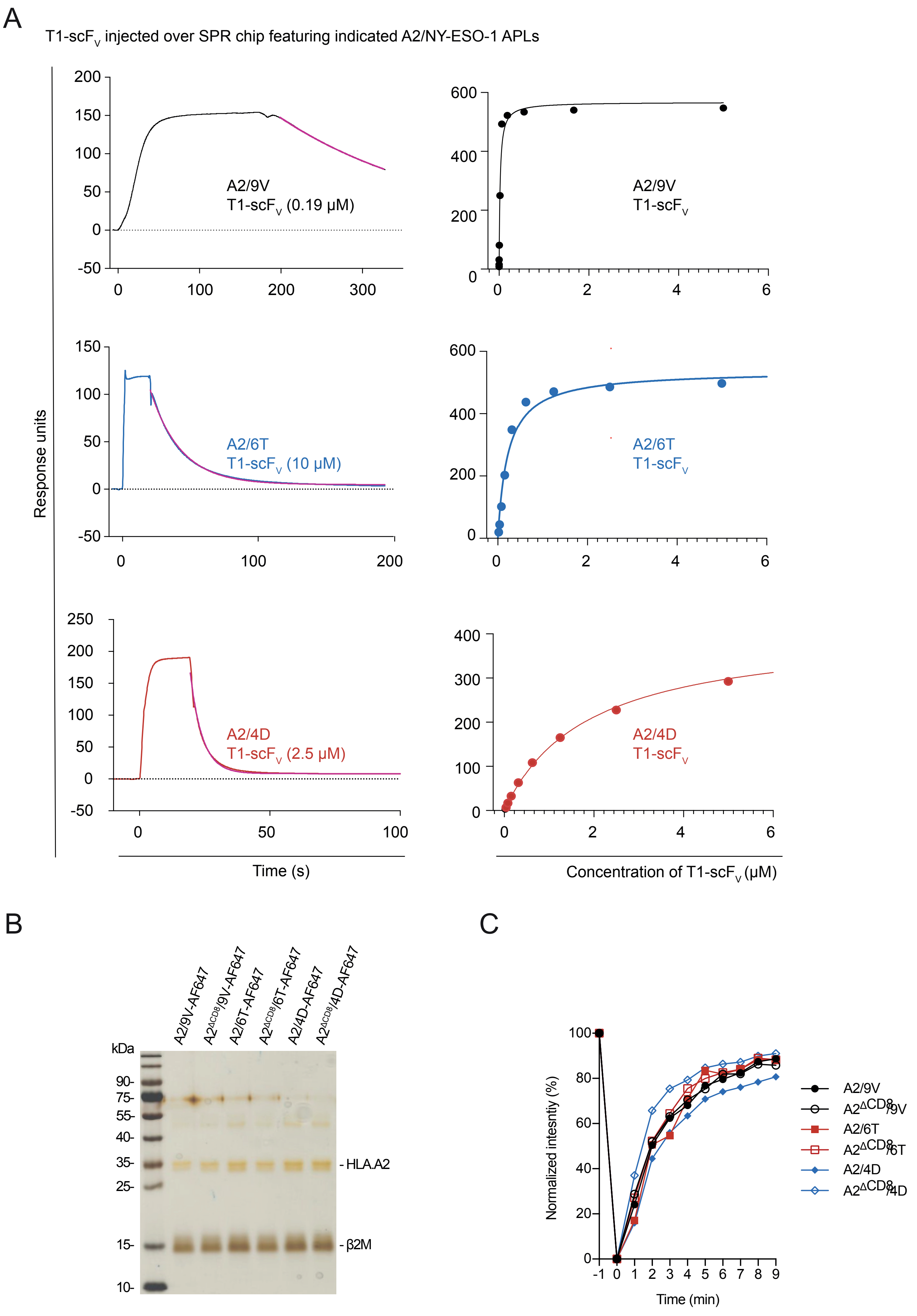

### Extended Data Figure 4

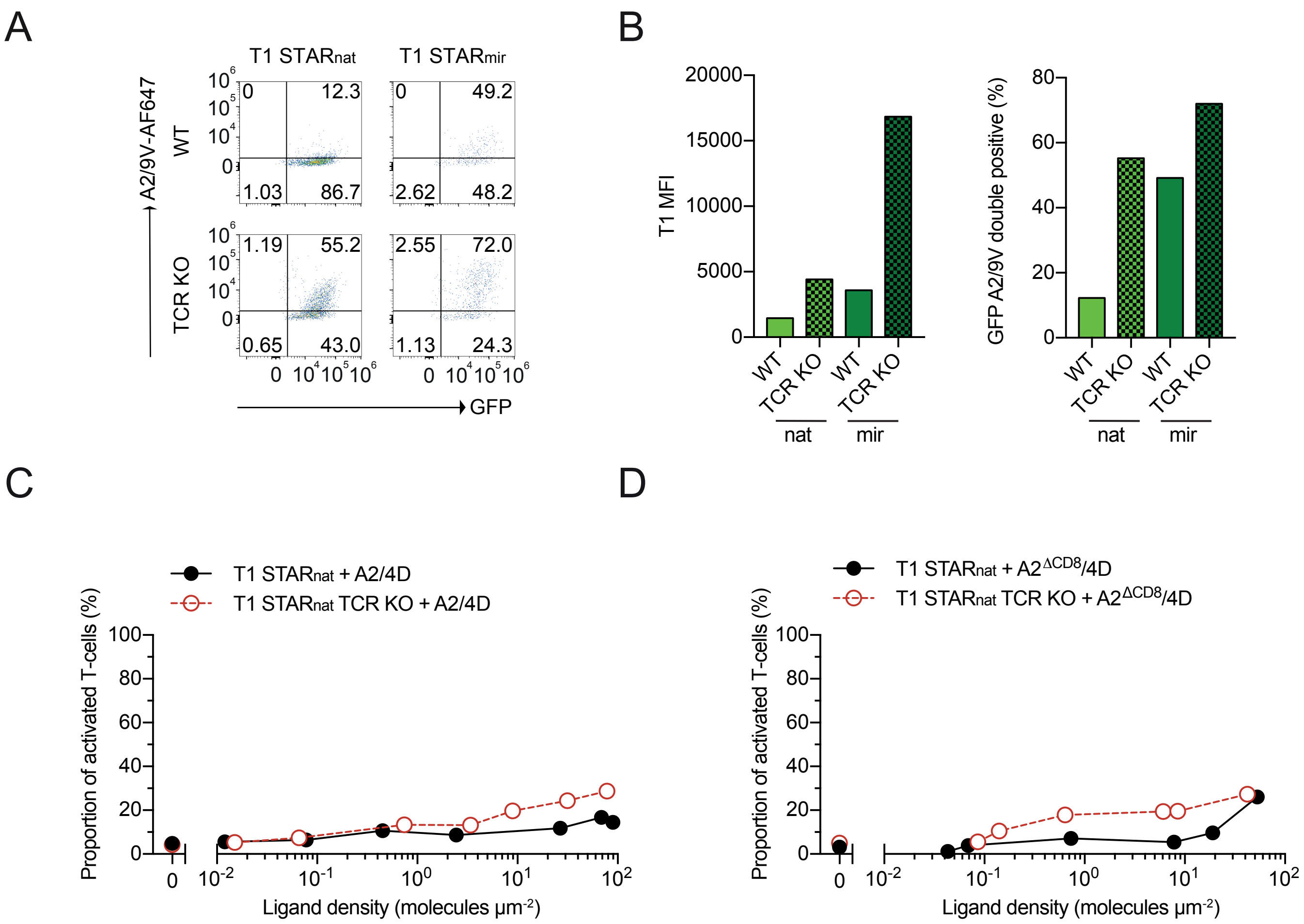

### Extended Data Figure 5

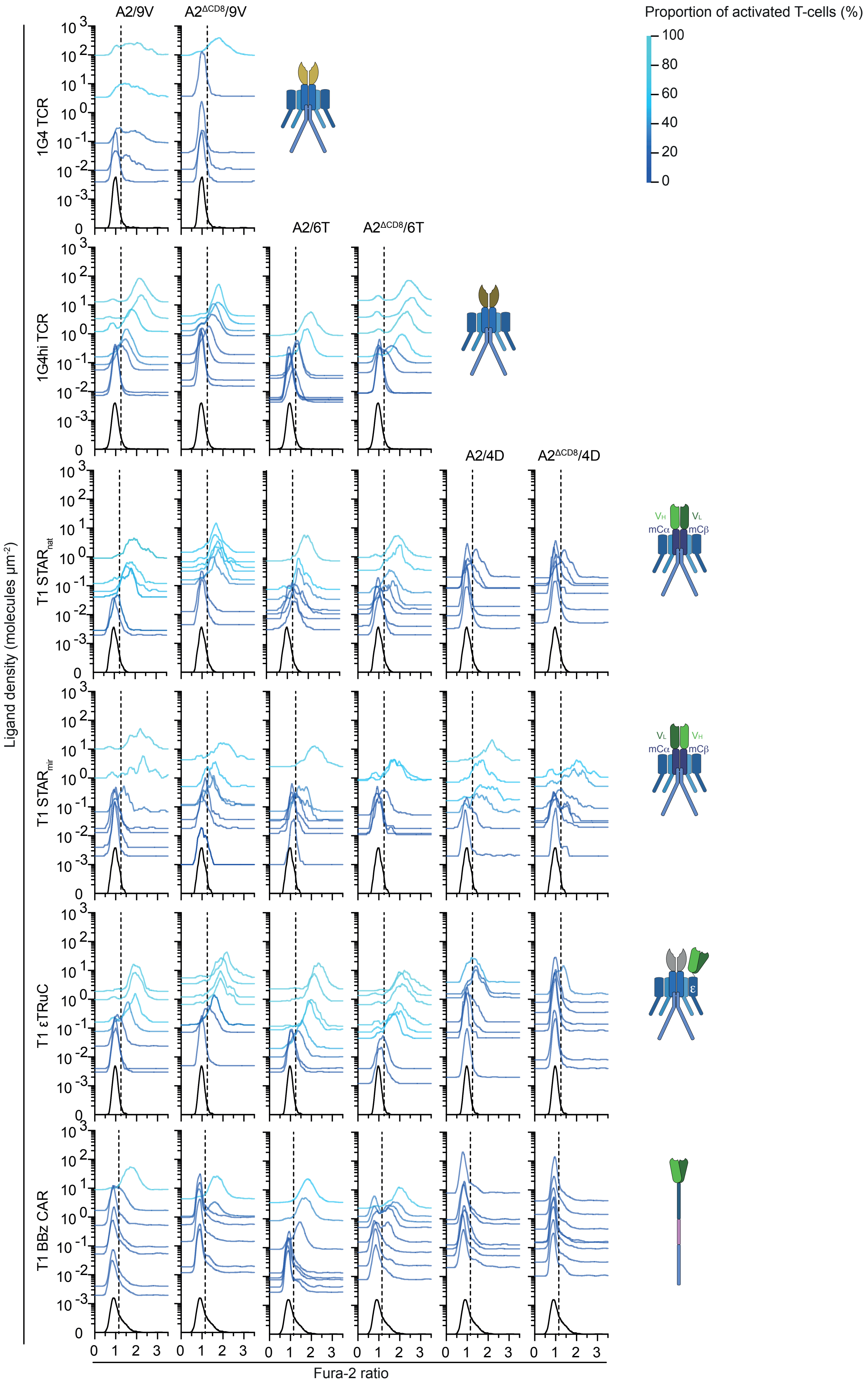

### Extended Data Figure 6

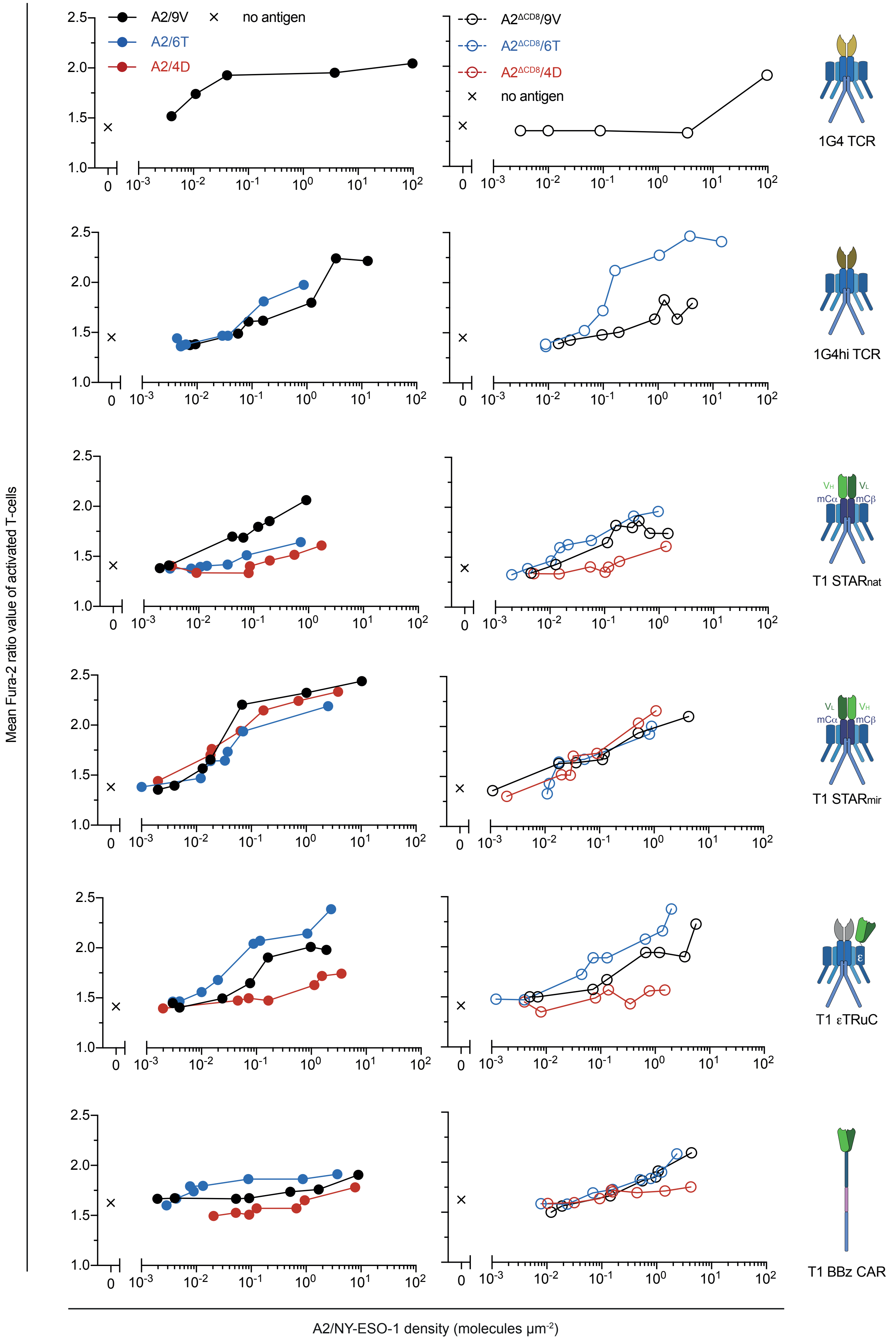

### Extended Figure 7

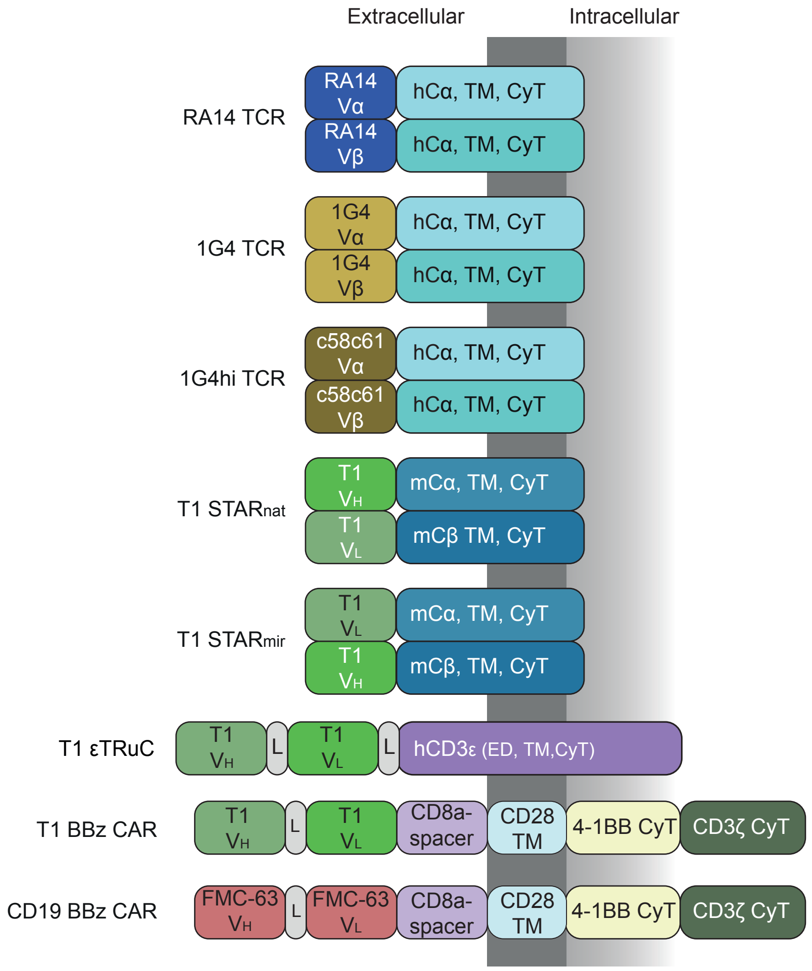
